# Spatiotemporal Dynamics of the Dihydrolipoyl Dehydrogenase LpdA Determine the Intrinsic Autofluorescence in Filamentous Actinobacteria

**DOI:** 10.1101/2025.03.03.640617

**Authors:** Denis Iliasov, Ricardo Santander Gualdron, Pilar Lörzing, Sandra Maaß, Buse Cinar, Bastian Wollenhaupt, Dietrich Kohlheyer, Dörte Becher, Christoph Loderer, Natalia Tschowri, Michael Schlierf, Vasily Zaburdaev, Thorsten Mascher

## Abstract

The hyphal nature of filamentous streptomycetes poses unique challenges to their multicellular lifestyle, since it requires organizing cellular functions at the scale of hundreds of micrometers length. Streptomycetes exhibit a strong and patchy autofluorescence of so far unknown origin in their hyphae that – as we demonstrate – is a natural property of filamentous actinobacteria. The foci are dynamic and evenly distributed throughout the hyphae, including the hyphal tips, where they are cell membrane-associated. Here, we resolved the high spatiotemporal dynamics of these foci during spore germination and vegetative growth in *Streptomyces venezuelae*. We isolated a fluorescent protein band and identified the responsible protein as dihydrolipoyl dehydrogenase LpdA. An *lpdA* deletion mutant lacked these fluorescent foci and showed minor deficiencies in growth and development. Heterologous LpdA production in *E. coli* and characterization of the enzyme verified that a flavin cofactor is responsible for the green autofluorescence. LpdA is highly conserved in actinomycetes as a part of multienzyme complexes involved in central metabolism. Such delocalized metabolic centers provide a potential solution to mycelial multicellular lifestyle, where diffusion becomes a major challenge.

## Introduction

Autofluorescence, or native fluorescence, is a natural cellular property that arises from intrinsic biomolecules acting as endogenous fluorophores. These molecules encompass a diverse array of fluorescent cellular structural components and metabolites, including flavins, nicotinamide adenine dinucleotide (NAD^+^), aromatic amino acids, and quinones, all of which exhibit fluorescence across the visible light spectrum.^1,2^ Thus, the fluorescence spectra of some NAD- and most flavin-containing enzymes usually overlap with the emission wavelengths of green fluorescent proteins (e.g., GFP), which are routinely employed to analyze protein expression and localization.^3^ However, autofluorescence is often characterized by a low signal-to-noise ratio and limited detection sensitivity.^1^ While autofluorescent molecules in eukaryotic cells (e.g., chlorophyll and lignin) can also exhibit intracellular localization due to the formation of cellular compartments, bacterial autofluorescence is usually distributed throughout the cell.^4^

Streptomycetes exhibit a significant and patchy autofluorescence that, hampers the characterization of low-abundance proteins in time and space by fluorescence microscopy employing for instance GFP-tagging.^5^ The nature of this hyphal green autofluorescent foci as a natural property of *Actinomycetales* has remained unexplored.^6,7,8^

The phylum *Actinomycetales* encompasses heterotrophic and chemoautotrophic bacteria, most of which are aerobic.^9^ *Streptomyces* sp. and other filamentous actinomycetes represent a group of soil- and marine-dwelling Gram-positive bacteria with several unique traits within the bacterial kingdom.^10^ These sporulating bacteria are characterized by filamentous growth into a multicellular mycelium.^11^ The life cycle of streptomycetes begins with spore germination, which is characterized by the formation of germ tubes at the poles.^12^ Subsequently, the germinated spores elongate into branching filamentous hyphae through apical growth by the polarisome.^13^ Growth of streptomyces by tip extension results in the formation of a multicellular mycelium without cell fission, thereby generating a cytoplasmic continuum that can easily stretch over hundreds of micrometers.^14^ The hyphae, however, are incompletely separated and compartmentalized through vegetative partitions, each containing multiple copies of the chromosome.^15^ With the onset of adverse conditions such as nutrient depletion, the vegetative mycelium undergoes cellular differentiation into reproductive structures called aerial hyphae. Each aerial hypha subsequently differentiates into chains of individual spores through coordinated chromosome distribution and controlled cellular division.^16,17,18^ Following differentiation, thick-walled, dormant spores are released that can survive unfavorable conditions for extended periods.^9^

In this study, we report on the spatiotemporal dynamics of the green autofluorescent foci as a natural property of vegetative mycelia in filamentous actinobacteria, including their localization during spore germination and vegetative growth in *Streptomyces venezuelae* and other actinomycetes. We reveal the autofluorescent component of these foci as the flavin-containing dihydrolipoyl dehydrogenase LpdA, an enzyme involved in central energy metabolism. Our experimental data, together with phenomenological theoretical model, indicate that the dynamic foci provide a means for spatial metabolism to ensure a balanced provision of central metabolites and energy equivalents along the growing hyphae in filamentous bacteria.

## Results

### Autofluorescence as a natural property of actinomycetes

While autofluorescence had been observed in streptomycetes and hampered the use of GFP-tagging for protein localization, the origin of this phenomenon has so far been elusive. Therefore, comprehensive fluorescence microscopy studies were performed on two model organisms of the genus *Streptomyces* (*S. venezuelae* (Fig. 1A) and *S. coelicolor* (Fig. 1B)), as well as *Actinomadura coerulea*, and two natural *Streptomyces* sp. isolated from the environment (Figure S1), in order to gain insight into this natural property of filamentous actinomycetes.

**Fig. 1.**
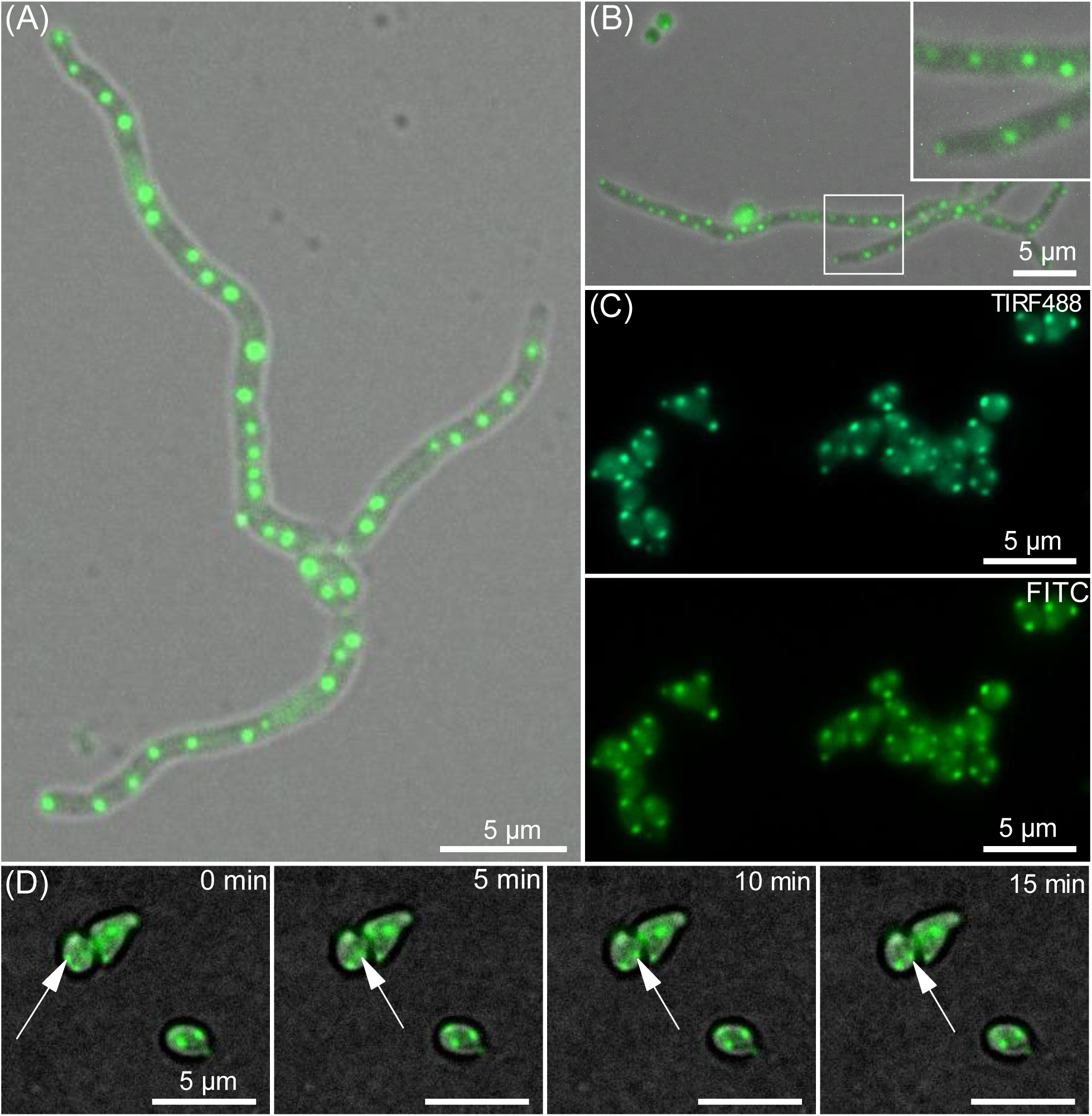
Autofluorescent imaging of *Streptomyces* species. **(A) *S. venezuelae*** and **(B) *S. coelicolor*** brightfield image overlaid with a fluorescence microscopy image showing hyphae with distinct, regularly spaced green autofluorescent foci. **(C)** TIRF microscopy on germinating *S. venezuelae* spores showing distinct foci with 488 nm (top) and 495 nm excitation (bottom). **(D)** TIRF microscopy on germinating *S. venezuelae* spores under continuous time-lapse imaging. White arrows indicate a dynamic foci moving inside the spores. Scale bars indicate 5 μm. For the corresponding movies, see Supporting Information Movies S1-S5.

All strains exhibited autofluorescence in the green light spectrum (490-575 nm) with excitation at 480 nm, with fluorescent foci regularly appearing in the hyphae as well as localized at tips and branching points. These latter spots correlate with the localization of the polarisome in streptomycetes.^13^ Remarkably, autofluorescent foci were also observed in the spores of *S. venezuelae* (Fig. 1C). Quantitative analysis of these foci consisted of measuring fluorescence intensity profiles along the growing hyphae to the hyphal ends and in comparison to the hyphal background autofluorescence, the foci showed a 2.5- to 4-fold higher autofluorescence (Figure S2).

Localization of the hyphal foci was determined by total internal reflection fluorescence microscopy (TIRFM) conducted on germinating spores (Fig. 1C and Figure S4). TIRFM enables the selective imaging of interfacial and near-surface processes (less than 200 nm) by eliminating background noise with high surface sensitivity.^19^ TIRFM of *Streptomyces* spores revealed association of the foci with spore poles and hyphal tips (Fig. 1C). These results suggest that the green autofluorescent foci on the hyphal tips are associated with the bacterial plasma membrane. Moreover, the foci exhibited remarkable stability on the hyphal tips during the germination of the spores and through vegetative growth (Fig. 1C, Figure S4, and Movie S5). In contrast, the pre-germination stage is distinguished by the movement of the foci in the spores (Fig. 1 (D) and Movie S5). The spatiotemporal dynamics, stability, and wide conservation in filamentous actinobacteria (Figure S1) raised the question of the molecular nature of these foci.

Fluorescence Lifetime Imaging Microscopy (FLIM) was performed on hyphae of *Streptomyces venezuelae* to elucidate the molecular characteristics of the autofluorescent foci. The intrinsic cellular fluorescence was excited with short pulses at 488 nm, and emission was collected between 500 and 600 nm. Both a weak background fluorescence signal in the cytosol and highly fluorescent foci could be observed (Figure S7 A and B), in agreement with previous intensity imaging. Within each foci, we found multiple, fluorescence lifetimes, indicating multiple components or states of the auto-fluorescent components (Figure S7 B). Fluorescence lifetime analysis identified three primary states, τ_1_=0.3 ± 0.1 ns, τ_2_=1.5 ± 0.3 ns, and τ_3_=4.7 ± 0.4 ns (Table S1). The mean intensity-weighted lifetime was <τ> int = 3.6 ± 0.1 ns (Figure S7 B and Table S1). The cytosolic background fluorescence was dominated by a fluorescent species with <τ> int = 0.8 ± 0.4 ns, which was significantly different from that of the foci. Using spinning disk confocal microscopy, we could demonstrate that the hyphal distribution of the green autofluorescent foci was uniform in all Z-planes (∼5 µm height) (Figure S4 and Movie S1 and S2). The lack of focus in the hyphae can be explained by the loss of the focus or by the ageing of the hyphae. Bacterial hyphae range in diameter from 0.8 to 1 µm, and foci were identified at a height of 1.8 µm (green; image 9 in Z-stack) with adjacent foci discernible at 2.4 µm (light blue; image 12 in Z-stack). Given the concurrence between measured Z-distance between neighboring detected foci and hyphal diameter, these findings suggest that the fluorescent molecules are evenly distributed in the Z-planes of the hyphae themselves.

### Isolation and identification of LpdA as a key component of fluorescent foci

The intracellular localization of the green fluorescent signals during vegetative growth and their observed dynamics during spore germination suggest a protein-associated origin of the foci. We therefore next aimed at isolating proteins showing autofluorescence from *S. venezuelae* to identify the origin of the observed foci. 15 µg of whole cell proteins isolated from *S. venezuelae* grown for 20 h or 48 h were separated using SDS-PAGE. As a control, the protein lysate was also subjected to 95 °C incubation for 10 min to ensure protein unfolding. Protein fluorescence was subsequently analyzed using a Typhoon multi-mode laser imaging system equipped with the AlexaFluor488 filters (Fig. 2B).

**Fig. 2.**
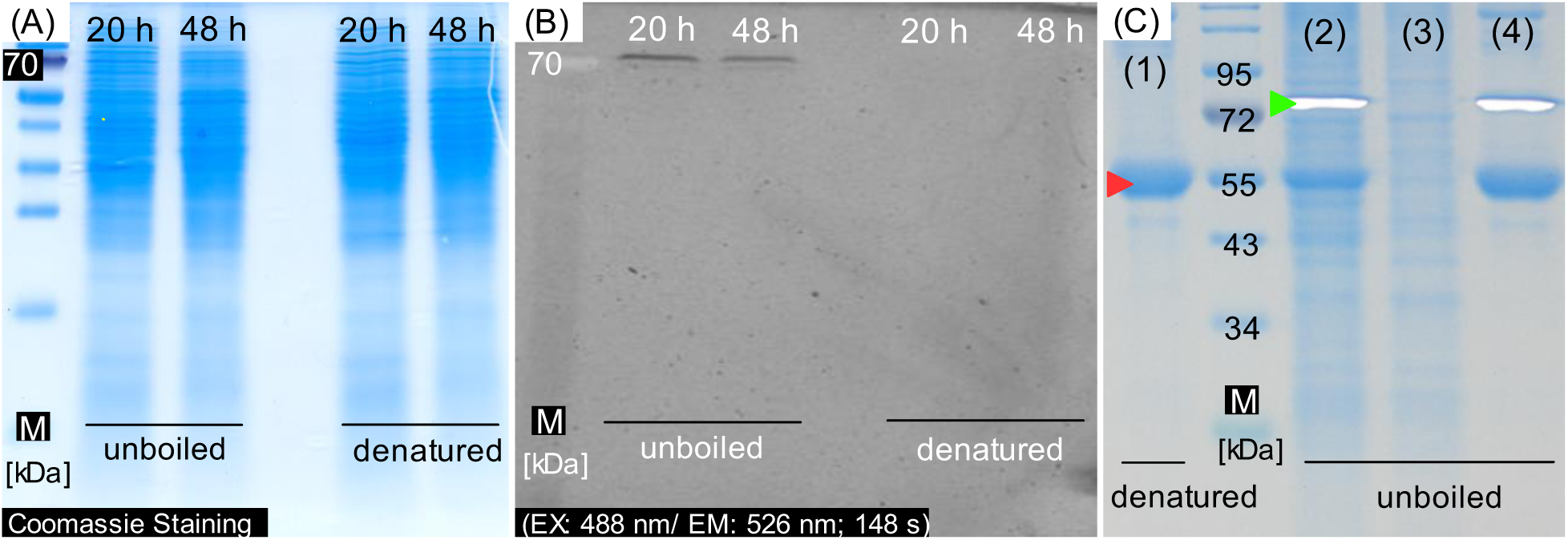
Autofluorescence of the *S. venezuelae* proteins (A) and (B). Proteins were isolated from *S. venzuelae* after cultivation at 28 °C for 20 h and 48 h. 15 µg of the isolated proteins per lane (in both undenatured and denatured states) were used for SDS-PAGE gel. Thermo PageRuler™ prestained protein ladder was used as marker (M). Fluorescence of the isolated proteins was detected using a Typhoon multi-mode laser imaging system (B) equipped with the AlexaFluor488 filters (PMT: 650; Filter: 555 BP 20 R6G, HEX, AlexaFluor532, exposure time of 148 s). For colorimetric visualization, staining was performed with Coomassie brilliant blue R-250 (left). **SDS-PAGE of heterologously expressed LpdA-6×His (C)**. M: Protein molecular weight marker. Lanes 1 and 4: Purified LpdA-6×His after denaturation (1) and without boiling form (4). Lane 2: Unboiled, total protein extract after expression of *E. coli* BL21 pET28b-LpdA-6×His. Lane 3: Unboiled, total protein extract of *E. coli* BL21. Arrows mark monomers (red) and homodimers (green) of LpdA. 15 µg of the isolated proteins per lane were used for SDS-PAGE gel. Thermo PageRuler™ prestained protein ladder was used as marker (M). Protein fluorescence was detected using a UV transilluminator. For colorimetric visualization, staining was performed with Coomassie brilliant blue R-250.

Green autofluorescence was only visible in the unboiled protein sample, with a single fluorescent band corresponding to an apparent molecular weight (MW) of 70 kDa. The heat denaturation step resulted in a loss of this fluorescent band, suggesting that the detected fluorescence is protein-associated. For identification of the corresponding protein, the fluorescent band of MW_app_ ∼70 kDa was excised from the gel and the proteins from within this gel sample were subsequently subjected to analysis using mass spectrometry. A total of 494 proteins were identified in this sample (Table S3). Protein candidates potentially responsible for foci formation were selected based on their abundance and their postulated fluorescent properties. The protein with the highest abundance was identified as a dihydrolipoyl dehydrogenase (LpdA, WP_041662298.1, 55 kDa). In its homodimeric form (110 kDa), this enzyme coordinates a flavin adenine dinucleotide (FAD) cofactor and constitutes a component of the multienzymatic complexes involved in primary metabolism.^20–22^ One of the characteristic properties of flavoproteins, including LpdA, is their autofluorescence in the green light spectrum (emission wavelength at 510 - 540 nm).^23,24^ These features and its overall abundance within the excised size range made LpdA a very promising candidate for the observed autofluorescence foci. We therefore generated and analyzed an *S. venezuelae lpdA* mutant.

The gene *lpdA* was deleted from the chromosome of *S. venezuelae* to determine the role of LpdA in the formation of the observed foci. Compared to the wild type strain, the resulting isogenic *lpdA* mutant showed an aberrant colony morphology (Fig. 3A), an inability to form pellicles (biofilms at the liquid-air-interface; Fig. 3C), and a slight growth defect (Fig. 3E). Microscopically, the *lpdA* deletion, while not affecting the diffuse intracellular autofluorescence level (Fig. 3 and Figure S2), led to a strain lacking visible green autofluorescent foci in the hyphae (Fig. 3), thereby demonstrating that LpdA is crucial for fluorescent foci formation.

**Fig. 3.**
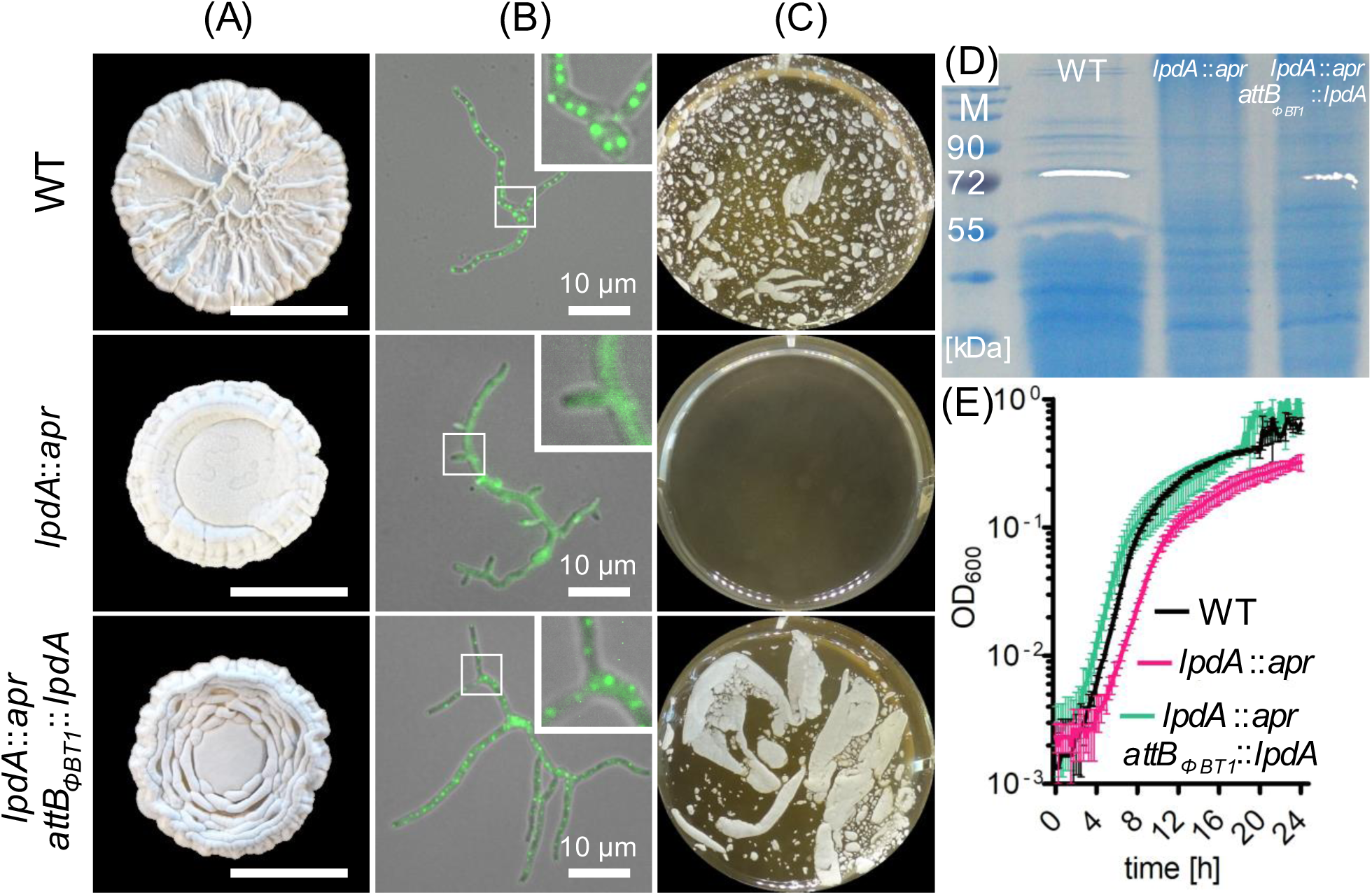
Phenotypes of *S. venezuelae* (WT), *lpdA* mutant (*lpdA*::*apr*), and the complemented *lpdA* mutant (*lpdA*::*apr*; *attBΦBT1*::*lpdA*). (A) Colony morphology of the *S. venezuelae* strains is shown for day 10 of growth on MYM agar. Scale bars indicate 1 cm. **(B)** Brightfield images overlaid with fluorescence microscopy images of the hyphae. Inset: Zoomed fraction of the full-sized image. **(C)** Ability to form pellicles (biofilms at the liquid-air-interface) of *S. venezuelae* strains during incubation in MYM medium**. (D)** SDS-PAGE of proteins from *S. venezuelae* and *lpdA* mutants. Proteins were isolated from *S. venzuelae* strains after cultivation at 28 °C for 20 h. Protein fluorescence was detected using a UV transilluminator shown as a white overlay. For colorimetric visualization, staining was performed with Coomassie brilliant blue R-25 (**E)** Growth curves (OD600) over 24 hours. Growth in MYM media of *lpdA* deletion strain (purple) and complemented strain (green) in comparison to the wild type (black).

The mutant phenotype was complemented by reintegration of *lpdA* under the control of its native promoter with pDI2-plasmid and expression from the *Φ_BT1_* integration site. This *lpdA* complementation strain regained the ability of *S. venezuelae* to form pellicles (Fig. 3C), and fluorescent foci, which were observed at the tip, branching points, and along the hyphae similarly to the pattern observed in the wild type (Fig. 3B). Importantly, the resulting foci in the *lpdA* complementation strain exhibited also 2.5- to 4-fold increase in measured fluorescence intensity in comparison to the background autofluorescence, which correlates very well with the original observations made for the wild type strain (Figure S2). The complemented *lpdA* mutant also regained some of the morphological characteristics displayed in colonies of the wild type strain. During growth curves, no significant differences were observed between the complemented *lpdA* strain and the wild type (Fig. 3E).

Proteins were extracted from *S. venezuelae* wild type, Δ*lpdA,* and the complemented mutant expressing *lpdA* and subjected to SDS-PAGE without incubation at 95 °C (Fig. 3D). In the *lpdA* deletion strain, the green autofluorescent band at ∼ 70 kDa was absent, while the complemented *lpdA* mutant exhibited autofluorescence under UV light excitation similar to the wild type strain, further supporting our hypothesis that the native form of LpdA in *S. venezuelae* is responsible for green autofluorescent hyphal foci formation and the green autofluorescent band.

### Purification and molecular properties of LpdA

For analyzing the protein properties of LpdA, *lpdA* was heterologously overexpressed in *E. coli,* and the corresponding protein was purified as C- or N-terminally His-tagged versions. Migration of fully denatured protein on SDS-PAGE was consistent with the predicted MW (LpdA) ∼ 55 kDa (Fig. 2C, red arrows and Figure S5). Due to the previously observed fluorescence at a 70 kDa band corresponding to LpdA in total protein samples isolated from *S. venezuelae* (Fig. 3D), purified LpdA samples from *E. coli* were also subjected to SDS-PAGE without incubation at 95 °C as a temperature denaturation step. Coomassie-stained and fluorescent bands corresponding to the apparent MW (∼70 kDa) could indeed be identified (Fig. 2C, green arrow) in agreement with the total protein extract from *S. venezuelae*. The fluorescent bands do not align with the expected MW of a LpdA monomer or LpdA-homodimer (110 kDa), which is likely due to the preserved tertiary structure of the isolated enzymes. Furthermore, the unboiled *E. coli* extract and purified sample showed a non-fluorescent LpdA band at MW ∼ 55 kDa (Fig. 2C, (2), (4)), which could be attributed to partial denaturation during purification and SDS-PAGE and the dissociation of FAD. Noteworthy, no fluorescent bands were observed in the non-denatured protein samples from *E. coli* BL21 without the LpdA expression plasmid used as a control (Fig. 2C, (3)). These findings indicate that the observed fluorescence of the proteins from *S. venezuelae* can indeed be attributed to the natural properties of the isolated LpdA enzyme. Furthermore, the enzymatic activity of the isolated protein towards lipoic acid was investigated, with the two isolated His-tagged LpdA versions exhibiting similar specific activity (Figure S6 A). Therefore, both LpdA demonstrated specific activity in the NADH-dependent reduction of lipoamide acid at 0.23 units per milligram of protein, representing a 6.4-fold increase in activity compared to the activity of the enzyme in the absence of lipoamide acid as a substrate.

To gain insight into the fluorescent properties of the isolated enzyme, the fluorescence spectrum of purified LpdA was measured (Fig. 4 and Figure S6 B). As the hyphal foci and proteins isolated from *S. venezuelae* exhibit green autofluorescence the excitation spectrum of 50 µM C- and N-terminally 6×His-tagged proteins was measured at 520 nm EM. The measured spectrum correlated with common excitation maxima of other fluorescent FAD-binding proteins (flavoproteins) at 360 nm, and 450 nm.^25^ Following the spectrum measurement at 480 nm EX, the emission peak of 50 µM LpdA was determined to be situated between 510 nm and 540 nm (Fig. 4, Figure S6 B, blue).

**Fig. 4.**
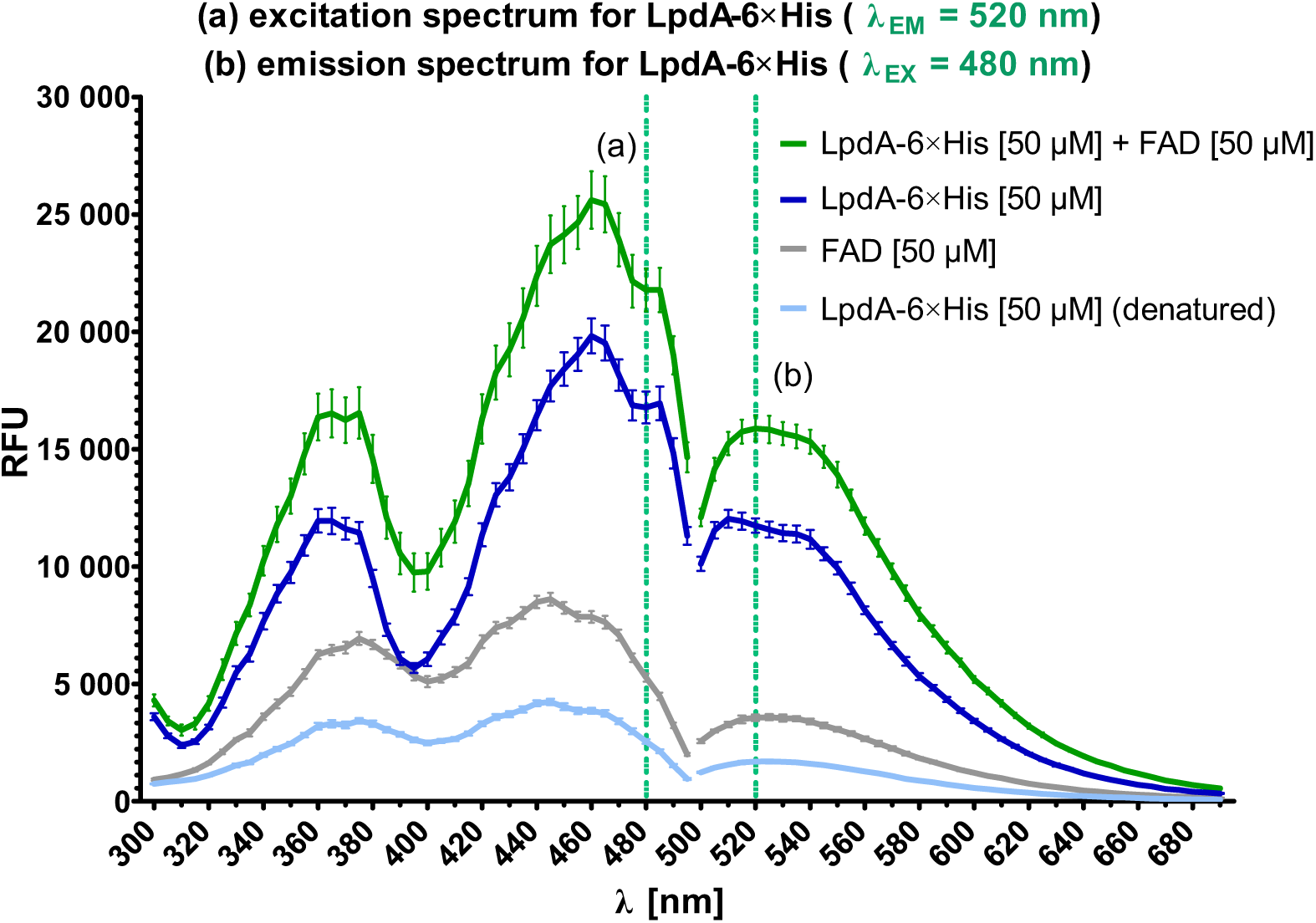
Fluorescence spectra of dihydrolipoyl dehydrogenase (LpdA-6×His). The excitation spectrum (a) was measured at 520 nm emission and the emission spectrum (b) was obtained at 480 nm excitation. Fluorescence spectra are shown for the purified 50 µM LpdA-6×His (blue), 50 µM FAD (gray), 50 µM denatured LpdA-6×His (light blue), and 50 µM LpdA-6×His after incubation with 50 µM FAD (green).

Denatured LpdA lacked visible peaks at 485 nm and 520 nm and exhibited a shift of the fluorescence spectrum to 450 nm and 530 nm (Fig. 4, Figure S6 B, light blue). These values correspond to the excitation and emission maxima of the flavin adenine dinucleotide (FAD) (Fig. 4, Figure S6 B, gray), an essential redox-active coenzyme for many metabolic enzymes. FAD therefore binds to the isolated dihydrolipoyl dehydrogenase. Heat-denaturation of the isolated enzyme resulted in a sevenfold reduction in the observed fluorescence (from 17,000 relative fluorescence units (RFU) to 2,500 RFU for excitation at 480 nm; from 12,000 to 1,700 for emission at 520 nm). This data indicates that the binding of FAD to LpdA may be the underlying cause of the intense green autofluorescence observed in the foci and isolated LpdA. Furthermore, the fluorescence spectrum of LpdA was quantified following incubation with FAD to ascertain the impact of the FAD-LpdA binding interaction on relative fluorescence. Strikingly, maximum LpdA fluorescence (EX at 488 nm and EM at 520 nm) increased by 5,000 RFU, indicating that the observed shift of fluorescence units can indeed be attributed to the FAD-bound LpdA enzyme, which may be responsible for the higher fluorescence of the foci compared to the background fluorescence.

Additionally, we measured the fluorescence lifetime of isolated LpdA in a fluorescence lifetime spectrometer. The fluorescence decay data was fitted by assuming a multi-exponential decay function, revealing a main lifetime component τ_1_ = 3.7 ± 0.13 ns, in good agreement with the lifetime *in situ* (Figure S7 C and Table S2). This further corroborates that autofluorescence in the foci in *S. venezuelae* cells likely originates from high concentrations of LpdA within the foci.

Phylogenetically, dihydrolipoyl dehydrogenases are a widely distributed group of enzymes with diverse metabolic functions.^26–29^ It is therefore surprising that autofluorescence foci can only be observed in filamentous actinobacteria. We therefore wondered whether the unique properties of these enzymes from this group of organisms are reflected by the conservation of their primary sequence. The amino acid sequence of LpdA from *S. venezuelae* shares strong homology to dihydrolipoyl dehydrogenases of both prokaryotes and eukaryotes. Sequence alignments indicate that all proteins exhibit high levels of conservation in the domains for FAD and NAD^+^ binding, a disulfide bridge, and a histidine-containing active site (Figure S8). Based on the alignment data, most selected enzymes exhibited identical FAD and NAD^+^ binding domains. The comparative analysis demonstrated higher similarity between all LpdA enzymes in actinobacteria and LpdA homologs in deltaproteobacteria than other LPDs. Although LpdA orthogonality was detected in actinomycetes, no phylum-specific conserved domains could be identified to explain the specific fluorescence in actinobacteria.

One possible explanation could be that only an actinobacterial-specific strong accumulation of LpdA homologs leads to the observed foci. This assumption points towards a physiological role of the spatiotemporal localization and distribution of the LpdA-containing foci, potentially to orchestrate the intracellular control of physiological processes in the long hyphae of *Actinomycetota*. We therefore aimed to investigate the cellular dynamics of foci formation and foci distribution in vegetative hyphae of *S. venezuelae*.

### Dynamic nature of the foci during spore germination and vegetative growth

Microfluidic single-cell cultivation under continuous time-lapse imaging was performed to capture the dynamic nature of the LpdA-containing foci during spore germination and hyphal vegetative growth in *S. venezuelae*and *S. coelicolor* (Fig. 5 and Figure S3, Movies S3 and S4).

**Fig. 5.**
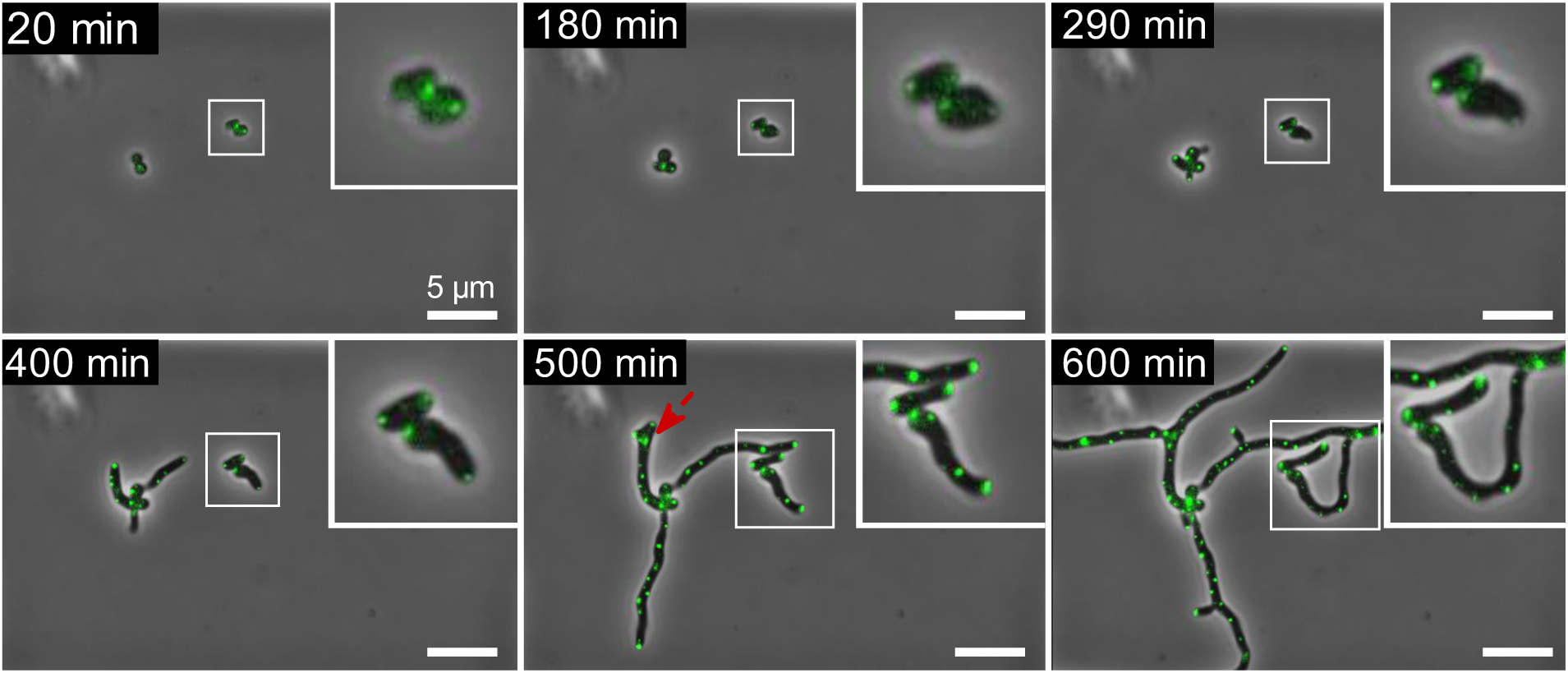
Spatiotemporal dynamics of LpdA-containing foci during vegetative growth of *S. venezuelae*. The strain was injected into the microfluidic single-cell cultivation system and cultivated at 28 °C for 12.5 h with phase contrast and fluorescence images taken in 10 min intervals. Red arrows indicate branching points. For the corresponding movies, see Supporting Information Movie S3.

Remarkably, the foci already exhibited movement in all spores during the period preceding germination (up to 150 min), indicating that the positions of the fluorescent LpdA-containing foci are not fixed within the spores. This period is also distinguished by the formation of new foci, which may be attributed to the activation of genes implicated in cell growth and primary metabolism.^30^ In the early germination phase, one green autofluorescent foci was directly associated with the hyphal tip (180 min) and exhibited remarkable stability throughout the vegetative growth. Several distinct lateral foci were also observed during vegetative hyphal growth. Images obtained during microfluidic cultivation confirmed that the foci accumulate at the branching positions of the hyphae (Fig. 5(A), red arrow), suggesting a potential role in both spore germination and vegetative hyphal growth.

Subsequently, a quantitative analysis was conducted to examine the spatiotemporal dynamics of the foci during vegetative growth (Figure S7 and S8). This demonstrated a direct correlation between the number of detected foci, the total fluorescence area, the general fluorescence intensity, and the active growth rate of the hyphae. The direct proportionality between hyphal growth and the increased number of foci over the entire cultivation period indicates that new foci are actively synthesized and are stable during vegetative growth.

However, no significant differences were observed between the fluorescence intensity and the average size of the foci formed over the entire cultivation period (Figure S7 and S8). These data indicate that foci synthesis occurs during spore germination and hyphal vegetative growth. Subsequently, the growth rate of the hyphae was calculated for the exponential growth phase, resulting in a value of 0.56 µm²/min for *S. venezuelae* and a value of 0.47 µm²/min for *S. coelicolor*. Additionally, the rate of foci synthesis for the two strains was determined to be approximately 0.85 foci/min, with 0.8 foci/min for *S. venezuelae* and 0.9 foci/min for *S. coelicolor*. As streptomycetes are characterized by apical growth at the hyphal tips, the values determined were normalized to the number of active tips (six for *S. coelicolor* and five for *S. venezuelae*) at the respective measurement time to calculate the tip-specific rates. The tip-specific growth rates were found to be 0.11 (µm²/min)/tip for *S. venezuelae* and 0.08 (µm²/min)/tip for *S. coelicolor,* while the tip-specific foci synthesis rates ranged from 0.15 to 0.16 (foci/min)/tip for both strains (supporting information S6). This suggests that the formation of hyphae is correlated with both the growth rate and number of active tips.

Random movements of the foci were also observed, which can be attributed to internal hyphal processes occurring during active growth. For instance, the movement of macromolecules towards the hyphal tip may be a contributing factor. Although a focus is associated with the hyphal tip from the moment of spore germination, laterally synthesized foci are characterized by a drift toward the growing hyphal tip. However, by characterizing the path of the selected foci, we observed that this drift is limited in distance, indicating a previously unknown spatiotemporal control of the localization and distribution of the foci in the hyphae (supporting information S8).

## Discussion

Native green autofluorescent foci are a unique natural characteristic of filamentous actinobacteria. In this work, we describe their localization and dynamics during spore germination and vegetative growth. We identified that the protein responsible for the autofluorescence in the foci is the dihydrolipoyl dehydrogenase LpdA, which was subsequently characterized regarding its fluorescent properties.

Dihydrolipoyl dehydrogenase (LpdA) belongs to the flavin-containing pyridine nucleotide-disulfide oxidoreductases, which play a significant role in primary cellular metabolism. LpdA is characterized by the presence of a reactive disulfide bridge in the active center and its involvement in catalysis of the NAD^+^-dependent oxidation of dihydrolipoyl groups covalently bound to the dihydrolipoyl transferase core component of multienzyme complexes.^24,26,31^ All lipoamide dehydrogenases described are homo-dimeric enzymes with two active sites at the dimer interface. Each monomer consists of four domains. In *S. venezuelae* LpdA, the N-terminal domain binds FAD and is in contact with the NAD-binding domain (48-360 amino acids), the central domain, and the interface domain (380-488 amino acids). Protein denaturation leads to the loss of tertiary and secondary structure and, consequently, also a loss of FAD binding capabilities in flavoproteins. The absence of discernible green autofluorescence peaks at the eGFP maxima (EX at 485 nm, and EM at 520 nm) for denatured samples indicates that the fluorescence spectra of isolated LpdA can be attributed to FAD binding and that the native structure of this enzyme is the sole source of the measured foci fluorescence (Fig. 4, and Figure S6 B). Furthermore, the fluorescence lifetimes of the detected foci and purified LpdA are attributable to the rigid binding of FAD molecules to LpdA.^32^

LpdA is an integral E3-subunit of three metabolic multienzyme complexes, including pyruvate dehydrogenase (PDH) complex, 2-oxoglutarate dehydrogenase (OGDH) complex, and branched-chain 2-oxo acid dehydrogenase complex.^24,33–36^ These complexes are composed of multiple decarboxylases or dehydrogenases (E1), dihydrolipoamide transacetylases (E2), and dihydrolipoyl dehydrogenases (E3) and function as catalysts for the utilization of sugars and amino acids.^37–39^ These multienzyme complexes of the α-keto acid dehydrogenase family act as catalysts for the oxidative decarboxylation of α-oxo acids and the associated production of reduction equivalents in the form of NADH. In contrast to the majority of microorganisms, in which these complexes are separated from each other, actinobacteria form the distinctive PDH/OGDH supercomplexes, which possess a dual function as PDC and OGDH and, as a result, constitute a crucial metabolic and multienzymatic complex within the glycolysis and TCA cycle of actinobacteria.^20,22,40,41^ Given that the synthesis of the apical cell wall in hyphae during vegetative growth in *Actinomycetales* is a highly energy-consuming process, the glucose level inside the spores increases due to the hydrogenation of trehalose.^12^ This process stimulates metabolic enzymes and increases the intracellular ATP concentration, thereby activating the polarisome and apical growth of hyphae.^13,42^ It was demonstrated that the genes encoding metabolic proteins were expressed within the initial 30 minutes following germination in *Streptomyces coelicolor.*^43^

In this study, LpdA-associated foci were also identified during microfluidic cultivation in the early germination phase inside spores of *S. venezuelae* and *S. coelicolor* (Fig. 1 and Figure S4; Movie S3 and S4). The reduced cellular differentiation and slowed growth observed in the *lpdA* deletion mutant can be attributed to the role of LpdA in primary metabolism. A comparable inhibitory impact of an *lpdA* knockout on growth was also observed in *E. coli*.^44^ Dihydrolipoyl dehydrogenase, as part of the branched-chain 2-oxo-acid dehydrogenase complex, is also involved in the catabolism of branched-chain amino acids and serves several metabolic functions in microorganisms. In the multicellular bacterium *Myxococcus*, these include the production of precursors for branched-chain fatty acids (BCFA) and precursors for morphogens that participate in cell-cell signaling.^24,45–47^ BCFA are involved in the synthesis of A-factor (2-isocapryloyl-3R-hydroxymethyl-γ-butyrolactone), which controls secondary metabolism and morphogenesis in *Streptomyces.*^48–50^ Previous research has demonstrated that the loss of branched-chain 2-oxo acid dehydrogenase complex enzymes results in strains exhibiting reduced morphological differentiation.^51,52^ The aforementioned data explain the reduced colony differentiation and the lack of pellicle formation during liquid cultivation of the *lpdA* mutant.

While most enzymes involved in bacterial central metabolism are distributed throughout the entire cytoplasm, the enzymes catalyzed the initial and final reactions of metabolic pathways can also have a specific localization in the cytoplasm.^53^ Furthermore, the existence of a unique PDH enzyme complex in actinobacteria with spatially defined localization has recently been discovered. All PDH complex and OGDH complex subunits are colocalized in *Corynebacterium glutamicum* at the cell poles, where they are associated with polar cell wall synthesis proteins.^21,54,55^ These findings are consistent with the colocalization found in our study of fluorescent LpdA-based foci with polarisome observed at the tips of hyphae. Moreover, additional fluorescent signals were observed in the mid-cell of larger *C. glutamicum* cells, which correlates with the even distribution of LpdA-associated foci seen in *S. venezuelae* hyphae.^21^ One potential explanation for this localization of hyphal foci as a part of metabolic multienzyme complexes is an interaction between one of the subunits and another protein. Nevertheless, no additional domains have been identified in the structure of multienzyme complexes that indicate an interaction with other proteins.^21,33^ Cumulatively, the even hyphal distribution of LpdA is potentially a common feature of actinomycetes. Although a high degree of sequence similarity was found between the FAD and NAD^+^ binding domains of dihydrolipoyl dehydrogenases (LPDs) from different organisms, these enzymes in actinomycetes exhibit significant divergence from other LPDs.

The intracellular localization and distribution of the detected native foci, whose fluorescence can be attributed to the properties of LpdA, and the role of this enzyme in metabolic processes inspired the hypothesis that the detected green autofluorescent foci in filamentous actinobacteria may potentially be due to the intracellular localization of unique metabolic multienzyme complexes. In light of this hypothesis, the localization of foci at the tips of hyphae, associated with the polarisome, and at branching points can be interpreted as correlating with sites of active growth. This hypothesis could also provide an explanation for the uniform distribution of foci in the hyphae. The observation of a defined spatial localization of an enzyme complex catalyzing two key central metabolism reactions alludes to the potential role of these green fluorescent foci in the intracellular availability of metabolic products (Fig. 6).

**Fig. 6.**
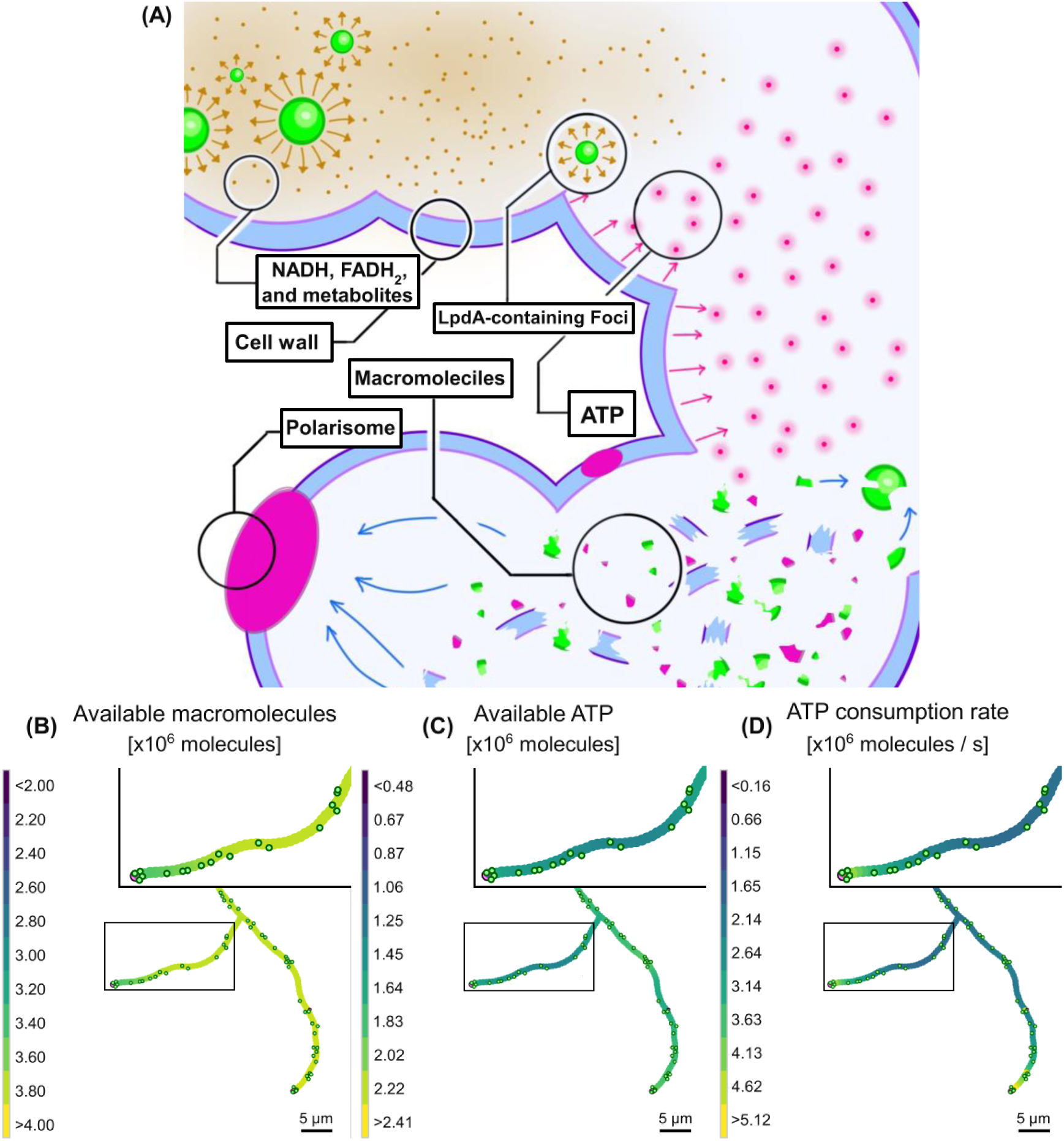
Agent based model showing the intracellular localization of green fluorescent LpdA foci and their potential role in the availability of metabolic products in the hyphae of actinomycetes. (A) Scheme of the components that were taken into account when building the model: cell wall, polarisome, LpdA-containing Foci, ATP, and macromolecules. Arrows indicate biochemical interactions between these components: production of NADH/FADH2, ATP production on the cell membrane, and cell envelope synthesis. Even though the figure shows NADH/FADH2 being released and diffused through the cytoplasm, we have simplified the model by assuming that these electron carriers get immediately consumed after they are released by the membrane-asociated foci. Once produced, ATP diffuses and gets consumed during the synthesis of macromolecules. The macromolecules diffuse until reaching a polarisome, where they are consumed to produce new cell components such as cell wall, among others**. (B)-(D) Growth simulation.** The concentration of freely diffusing macromolecules (B) and ATP concentration (C) are demonstrated to the time point of 4 h 48 min. The color scale represents the concentration of ATP or macromolecules in 10^6^ of molecules per cytoplasm segment. The panel (D) shows the rate at which ATP is consumed in each cytoplasm segment. The color scale represents the consumption rate in 10^6^ of molecules per second per cytoplasm segment. The LpdA-containing foci are represented by the green dots, and the polarisome are represented by the purple dots at the tips of the hyphae. Insets show magnified parts of the hyphae. See Supporting Information S7.

To corroborate this hypothesis, we developed an agent-based model for the growth and branching behavior of *Streptomyces* spp. as a function of LpdA-containing foci that takes into account the aspects and is illustrated in Fig. 6A. Here, the multicellular process of hyphae growth was simulated where the growing and moving foci serve as a source of metabolic products required for the biosynthesis of macromolecules and cell components. Remarkably, using the parameter values either known from the literature or deduced from our experimental data, the overall growth of the filamentous structure and its branching were recapitulated (Fig. 6 (B)-(D); the LpdA-containing foci as green dots; the polarisome as purple dots at the tips of the hyphae).

By incorporating cellular components including discrete foci as sites of localized ATP production, polarisome for tip extension, and continuous fields representing ATP and macromolecule gradients, the model successfully reproduces experimentally observed growth patterns and foci count. The model shows that *Streptomyces* hyphae likely exhibit spatial heterogeneity of metabolic activity along the filament. This heterogeneity arises from the interplay between localized ATP production at foci sites, random motion of foci, and elevated consumption near growing tips, where energy-intensive processes like cell wall synthesis occur. The model’s ability to reproduce both normal growth patterns and phenotypic variations like hyperbranching by parameter adjustments demonstrates its usefulness for studying the impact of perturbations on the multicellular organization of filamentous actinobacteria. This model illustrates how the localized condensation of LpdA in distinct dynamic aggregates in concert with polarisome can drive the multicellular growth of the *Streptomyces* bacteria.

Taken together, it is tempting to suggest that the dynamic localization and distribution of LpdA-containing foci highlight the spatial distribution of metabolic multienzyme complexes, as previously shown for *C. glutamicum*. In filamentous actinobacteria this need for a spatiotemporal control of central metabolism is emphasized by the hyphael and multicellular nature of the vegetative mycelium. This situation is reminiscent of the identical challenge faced by fungal mycelium and stands in contrast to most bacteria, in which diffusion of metabolites is not limiting due to the small size of the individual cells. Future research on the formation and composition of the foci-associated metabolic enzyme complexes is needed to further determine the coordination of spatiotemporal metabolism in *Actinomycetales*.

## Supporting information

supplementary figurs

S6_MSCC_Data

S7_Agent_based_model

S8_Theory_for_agent_based_model

## Acknowledgments

The authors thank Sebastian Grund (Greifswald, Germany) and Katrin Gunka (Hannover, Germany) for technical support.

## Materials and methods

### Bacterial strains, plasmids, and oligonucleotides

Strains and plasmids used are shown in Table 1, and oligonucleotides used are shown in Table 2.

**Table 1.** Strains and plasmids used in this study.

| Strain | Relevant genotype/comments | Source or reference |
| --- | --- | --- |
| ***Actinomadura* sp.** | Natural isolate | Isolated from dried *S. suricatta* samples |
| ***E. coli*** |  |  |
| ET12567/pUZ8002 | *dam*, *dcm*, *hsd*, Kan<sup>R</sup>, Cm<sup>R</sup> | 56 |
| BW25113/pIJ790 | $\Delta(araD-araB)567$, $\Delta lacZ4787(::rrnB-4)$, *lacI*p-4000(*lacI*<sup>Q</sup>), λ-, *rpoS*369(Am), *rph*-1, $\Delta(rhaD-rhaB)568$, *hsdR*514, Cm<sup>R</sup> | 57 |
| DH10β | F-mcrA $\Delta(mrr-hsdRMS-mcrBC)$<br>$\Phi 80dlacZ\Delta M15 \Delta lacX74 endA1 recA1$<br>*deoR* $\Delta(ara, leu)7697$ *araD*139 *galU*<br>*galK nupG rpsL* λ- | Strain Collection |
| BL21(DE3) | F- *ompT* *hsdS*<sub>B</sub> (*r*<sub>B</sub>-, *m*<sub>B</sub>-) *gal dcm*<br>(DE3) | Novagen |
| TME4835 | DH10β pIJ773, Apr<sup>R</sup> | |
| TME4841 | DH10β 3E09, Kan<sup>R</sup>, Amp<sup>R</sup> | This study |
| TME4846 | BW25113 pIJ790, Cm<sup>R</sup>;<br>3E09, Kan<sup>R</sup>, Amp<sup>R</sup> | This study |
| TME4852 | BW25113 pDI1 (3E09-*lpdA*:: *apr*, Apr<sup>R</sup>),<br>Kan<sup>R</sup>, Amp<sup>R</sup> | This study |
| TME4853 | ET12567 pUZ8002, Kan<sup>R</sup>, Cm<sup>R</sup>;<br>pDI1 (3E09-*lpdA*:: *apr*, Apr<sup>R</sup>), Kan<sup>R</sup>,<br>Amp<sup>R</sup> | This study |
| TME4857 | DH10β pDI2 (*lpdA*), Hyg<sup>R</sup> | This study |
| TME4858 | ET12567 pUZ8002, Kan<sup>R</sup>, Cm<sup>R</sup>; pDI2<br>(*lpdA*), Hyg<sup>R</sup> | This study |
| TME4861 | BL21(DE3) pDI3 (pET28b(+)-P<sub>T7</sub>-*lpdA*-6xHis), Kan<sup>R</sup> | This study |
| TME4862 | BL21(DE3) pDI4 (pET28b(+)-P<sub>T7</sub>-6xHis-*lpdA*), Kan<sup>R</sup> | This study |
| ***S. coelicolor* M600** | A plasmid free-derivative of the wild type strain | 58 |
| ***Streptomyces* sp.** | Natural isolate | Isolated from dried *V. salvator* samples |
| ***Streptomyces* sp.** | Natural isolate | Isolated from dried *P. panini* samples |
| ***S. venezuelae*** |  |  |
| NRRL B-65442 | Wild type | 59 |
| TMS0210 | NRRL B-65442 *lpdA::apr*, Apr<sup>R</sup> | This study |
| TMS0211 | NRRL B-65442 *lpdA::apr*, Apr<sup>R</sup>; *attB<sub>ΦBT1</sub>::pDI2* (*lpdA*), Hyg<sup>R</sup> | This study |
| **Plasmids** |  |  |
| pIJ773 | Plasmid template for amplification of the *apr-oriT* cassette for 'Redirect' PCR-targeting | 60 |
| pIJ790 | Modified λ RED recombination plasmid [*oriR101*] [*repA101* (ts)] *araBp-gam-be-exo*, Cm<sup>R</sup> | 60 |
| pUZ8002 | RP4 derivative with defective oriT, Kan<sup>R</sup> | 56 |
| 3E09 | SuperCos1 derivative carrying the *lpdA* region from *S. venezuelae*; Kan<sup>R</sup>, Amp<sup>R</sup> | http://strepdb.streptomyces.org.uk |
| pDI1 | 3E09 (*lpdA::apr*, Apr<sup>R</sup>), Kan<sup>R</sup>, Amp<sup>R</sup> | This study |
| pSS172 | pIJ10770 carrying *mcherry* for the construction of C-terminal fluorescent gene fusions, Hyg<sup>R</sup> | 61 |
| pDI2 | pIJ10770-*lpdA*, Hyg<sup>R</sup> | This study |
| pET28b(+) | T7lac promoter, Kan<sup>R</sup>, pBR322 ori | Novagen |
| pDI3 | pET28b(+)-P<sub>T7</sub>-*lpdA*-6xHis, Kan<sup>R</sup> | This study |
| pDI4 | pET28b(+)-P<sub>T7</sub>-6xHis-*lpdA*, Kan<sup>R</sup> | This study |

**Table 2.**
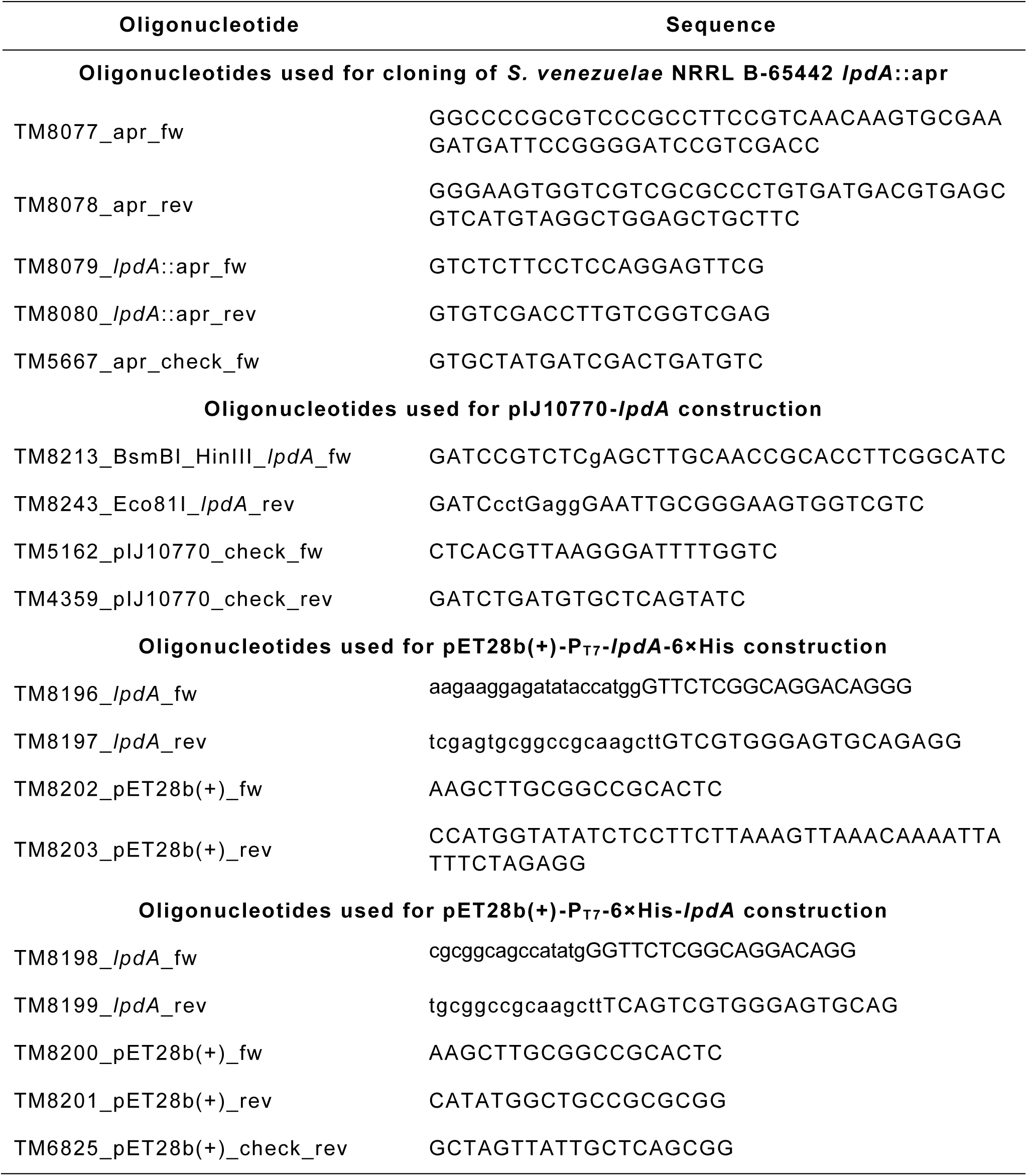
Oligonucleotides used in this study.

### Bacterial strains, growth conditions, and conjugations

All *Escherichia coli* strains used in this study (Table 1) were routinely grown in Lysogeny broth (LB-Medium (Luria/Miller), Carl Roth GmbH + Co., KG, Karlsruhe, Germany) at 37 °C or 28 °C with agitation. When required, the following antibiotics were added to the growth medium: 100 μg⋅mL^-1^ ampicillin (Amp^R^), 50 μg⋅mL^-1^ kanamycin (Kan^R^), 100 μg⋅mL^-1^ hygromycin (Hyg^R^), 50 μg⋅mL^-1^ apramycin (Apr^R^), or 25 μg⋅mL^-1^ chloramphenicol (Cm^R^). *E. coli* DH10β was used for plasmid and cosmid propagation, and BL21 (DE3) for protein overexpression. *E. coli* BW25113^57^ containing a λ RED plasmid, pIJ790, was used to create the *lpdA* (*vnz_09030*; *SVEN_1842*) disruption 3E09-cosmid (pDI1). Conjugation between *S. venezuelae* NRRL B-65442 and *E. coli* strains ET12567 containing pUZ8002^56^ were performed as described by Bibb *et al*.^62^ *Streptomyces* strains and *Actionomadura* sp. used are summarized in Table 1 and were cultivated in maltose-yeast extract-malt extract (MYM) medium prepared with tap and deionized water (1:1) and supplemented with 200 mL of R2 trace element solution per 100 mL.^63^ Liquid cultures were incubated aerobically at 28 °C and 170 rpm. When required, MYM agar contained 50 μg⋅mL^-1^ kanamycin, 100 μg⋅mL^-1^ hygromycin, or 50 μg⋅mL^-1^ apramycin.

### Colony morphology

Spore suspensions of *Streptomyces* strains were spotted on MYM agar plates and incubated according to the general growth conditions. Pictures of colonies were taken on a black background using a P.CAM360 (1.48x magnification, overhead light level 3) [TU Dresden] and edited for light and contrast using the open-source software Fiji.^64^

### Growth assays

To determine viability of *S. venezuelae* strains in MYM medium, optical density values at the wavelength 600 nm (OD_600_) were measured using a plate reader Synergy™ HTX multi-mode microplate reader from BioTek (Winooski, USA) set at 28 °C and shaking at 700 rpm. 1.5 mL of MYM was pipetted into the wells of a 12-well cell culture plate (677102 Greiner Bio-One) and inoculated with 20 µL spore suspension (10^6^ spores⋅mL^-1^). OD_600_ measurements were recorded every 15 min for 30 h and data was subsequently visualized using GraphPad Prism (vers. 5, San Diego, California). Each experiment was performed in triplicate. To characterize the *S. venzuelea* strains with regard to their capacity to form pellicles (*i.e*., biofilms at the liquid-air interface), the spores from the strains were subjected to an incubation process. The spores were incubated in 1.5 mL MYM at 28°C for 24 hours, without shaking. Pictures of colonies were taken on a black background using a P.CAM360 (1.48x magnification, overhead light level 3) [TU Dresden] and edited for light and contrast using the open-source software Fiji.^64^

### Fluorescence microscopy

The *Streptomyces* stains and *Actinomadura* sp. were incubated in MYM medium for 18 h at 28 °C. Cells were applied to 1% (w/v) UltraPure agarose pads and fluorescence microscopy was performed using an Axio Observer 7 inverse microscope (Carl Zeiss, Jena, Germany) equipped with standard mCherry (EX:587/EM:610; exposure time of 1.5 s) and eGFP (EX:488/EM:509; exposure time of 5 s) filter sets. *Streptomyces* strains were incubated in MYM medium for 4 h and subsequently used for TIRF microscopy with a DeltaVision Ultimate Focus microscope (GE Healthcare Life Sciences, Chicago, IL, USA) equipped with standard FITC (EX:475/EM:528; exposure time of 3 s), mCherry (EX:575/EM:630; exposure time of 1.5 s) and TIRF lasers (488 nm). The acquired images were analyzed using the open-source platform Fiji.^64^ Microscopy of *Streptomyces* stains was performed in biological and technical triplicate.

### Sample preparation for microscopy

Glass slides and coverslips (150 µm thickness) were cleaned twice by sonicating for 10 min in 5% Mucasol, 10 min in ethanol and drying with nitrogen gas. An 1% agarose pad was prepared according to Skinner *et al*.^65^ A solidified agarose pad was placed on a glass slide, 2 µl cell suspension were placed on top and covered with a coverslip.

### Spinning disk confocal microscopy

Confocal imaging was performed using a Nikon Ti-E Spinning Disk microscope equipped with a Yokogawa CSU-X1 confocal scanner unit at 5,000 rpm scanning speed. A 100 × Apo TIRF objective (1.49 NA) with an additional 1.5 × tube lens was used and an Andor Ixon Ultra 888 EMCCD camera was used for detection. Cellular fluorescence was recorded with 488 nm laser excitation (56 mW before the objective) and emission light was filtered using a dual band filter 433/530 HC (Semrock). Images were acquired as z-stack with 0.2 µm step size and 400 ms exposure time for each frame.

### Fluorescence lifetime imaging microscopy

FLIM experiments were carried out using a Leica Stellaris 8 FALCON confocal FLIM microscope with LAS-X software. Samples were excited with 488 nm line of the white-light laser, and a repetition rate of 40 MHz. Photon arrival times were recorded with a HyD X 2 detector, covering the emission range between 500 – 600 nm. A 63× objective with additional 2.38 × zoom was used. Acquisition was performed through a 512 × 512 pixel image format with 0.151 µm pixel size at a speed of 400 Hz and a pinhole size of 1 airy unit. Images were recorded as 100 frames time average. Laser power was set between 5 to 10% to operate at count rates below 1 photon/pulse.

Image processing was done using LAS-X software. A 3-exponential tail fit was used for FLIM data analysis, having a time window of 20 ns. FLIM images were fitted with an intensity threshold of 10 counts to remove the background and represented as intensity weighted lifetime.

### Fluorescence lifetime measurements

The fluorescence decay curves were obtained with a FluoTime 300 fluorescence lifetime spectrometer (PicoQuant, Germany) by time correlated single photon counting. The instrument response function was obtained experimentally using Ludox as scattering standard. A pulsed laser light source (485 nm) at 11.4 MHz pulsed mode was used for excitation. Emission data were collected at 530 ± 13.5 nm. For the fluorescence decay curve a double-exponential decay function was assumed:

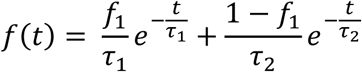

Where τ_1_ and τ_2_ are the lifetimes of the two components, f_1_ the fraction of the first component, and t the time. This biexponential function f(t) is then convolved with the IRF h(t) to account for the instrument response:

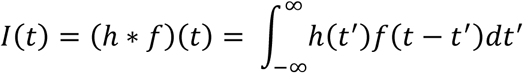

### Microfluidic cultivation conditions

Microfluidic single-cell cultivations (MSCCs) were performed using a previously described version of the polydimethylsiloxane (PDMS) cultivation device.^67^ The microfluidic cultivation system consists of eight arrays of monolayer cultivation chambers (length × width × height = 90 × 40 × 1 μm), with 40 chambers each. The fabrication of the chip was performed as described by Grünberger *et al*.^68,69^ Fluorescence microscopy and incubation of the microfluidic cultivation system was carried out using a motorized inverted fluorescence microscope (Nikon Eclipse Ti, Nikon, Japan) equipped with a 100× oil immersion objective (CFI Plan Apo Lambda DM 100X, NA 1.45, Nikon Instruments, Germany) and an incubator (PeCon Series 2000, PeCon GmbH, Germany) for temperature control. The microfluidic cultivation chips were inoculated with spore suspensions of *S. venezuaelae* NRRL B-65442 and *S. coelicolor* M600 (both 10^9^ spores⋅mL^-1^).^70^ During cultivation, a syringe pump (NeMESYS, Cetoni GmbH, Germany) was used to continuously supply the cells with fresh Gauze mineral medium (GAU) at a flow rate of 200 nL·min^-1^. Growth chambers most suitable for imaging were selected manually. Phase contrast and fluorescence time-lapse images of growing microcolonies were taken in 10 min intervals using an ANDOR LUCA R DL604 CCD camera. Fluorescence images were captured with an exposure time of 700 ms using a 300-watt xenon light source (Lamda DG-4, Sutter Instruments, USA) at 25% of maximum intensity and the appropriate optical filters (FITC (EX:470/EM:525); AHF Analysentechnik AG, Germany).

### Protein isolation and SDS-PAGE

MYM medium was inoculated with *S. venezuelae* spores to achieve a final concentration of approx. 10^6^ spores·mL^-1^ and incubated aerobically for 20 h and 48 h. Cells were harvested by centrifugation (4 °C, 10,000 g, 15 min) and washed once in ice-cold 20 mM Tris-HCl (pH 8). Following harvesting, the cell pellet was resuspended in 20 mM Tris-HCl (pH 8) with 1×EDTA-free protease inhibitors (Roche) and frozen in liquid nitrogen. Cells were subsequently lysed by sonication for 30 s at 60% power over 10 cycles, with 30 s on ice in between cycles. Cell debris was removed by centrifugation (4 °C, 13,000 g, 15 min). Protein concentration of cell lysates was determined using the Bradford assay and total protein concentration of each sample was adjusted to 1 mg⋅mL^-1^.

For the characterization of the isolated proteins regarding fluorescence, 15 µL of the samples were mixed with 5 µL 4×SDS loading dye (60 mM Tris-HCl (pH 6.8), 10% (v/v) glycerol, 2.5% (v/v) β-mercaptoethanol, 2% (w/v) sodium dodecyl sulfate, 12.5 mM ethylenediam-inetetraacetic acid, 0.025% (w/v) bromophenolblue) and run on a 12.5% SDS polyacryl-amide gel (180 V, 80 min) before and after 10 min incubation at 95 °C (for denaturation). Protein size was verified using either the Color Prestained Protein Standard (NEB; Ipswich, USA) or Thermo PageRuler™ prestained protein ladder (Waltham, MA, USA). Protein fluorescence was then detected using a Typhoon multi-mode laser imaging system (Filter: 555 BP 20 R6G, HEX, AlexaFluor532, exposure time of 148 s). For colorimetric visualization, staining was performed with Coomassie brilliant blue R-250 (SigmaAldrich, St. Louis, USA).

### Mass spectrometry

Gel lanes were excised and subjected to trypsin digestion as described earlier.^71^ For the subsequent LC-MS/MS measurements, the digests were separated by reversed phase column chromatography using an EASY nLC II (Thermo Fisher Scientific) with self-packed columns (OD 360 μm, ID 100 μm, length 20 cm) filled with 3 µm diameter C18 particles (Dr. Maisch, Ammerbuch-Entringen, Germany). Following loading/ desalting in 0.1% (v/v) acetic acid in water, the peptides were separated by applying a binary non-linear gradient from 1-99% (v/v) acetonitrile in 0.1% acetic acid over 77 min. The LC was coupled online to a LTQ Orbitrap mass spectrometer (Thermo Fisher, Bremen, Germany) with a spray voltage of 2.6 kV. After a survey scan in the Orbitrap (r = 30,000), MS/MS data were recorded for the five most intensive precursor ions in the linear ion trap. Singly charged ions were not considered for MS/MS analysis. The lock mass option was enabled throughout all analyses.

After mass spectrometric measurement, database search was performed using MaxQuant (version 2.3.1.0) with databases of *Streptomyces venezuelae* NRRL B-65442 (ATCC 10712) (obtained from NCBI on 23/08/31, txid953739, 24,650 entries).^72^ Common laboratory contaminants and reversed sequences were included by MaxQuant. Search parameters were set as follows: Trypsin/P specific digestion with up to two missed cleavages, methionine oxidation and N-terminal acetylation as variable modification, match between runs with default parameters enabled. The FDRs (false discovery rates) of protein and PSM (peptide spectrum match) levels were set to 0.01. Two identified unique peptides were required for protein identification. iBAQ values were calculated in MaxQuant with default settings as proxy for protein abundance.^73^

The MS proteomics data discussed in this publication have been deposited to the ProteomeXchange Consortium via the PRIDE partner repository^74^ with the dataset identifier PXD055355 (Reviewer account details: Username; Password, l7IsyDF3ZkF1).

### Construction and complementation of a *S. venezuelae lpdA* null mutant

An *lpdA* mutant strain was generated using the ‘Redirect’ PCR targeting method.^60^ The *lpdA* coding sequence (*vnz_09030*; *SVEN_1842*) on the cosmid vector 3E09 [http://strepdb.streptomyces.org.uk/] was replaced with a single apramycin-resistance (*apr*) cassette. Here, *E. coli* BW25113 containing pIJ790 were transformed with cosmid vector 3E09 and *lpdA* was replaced with the apramycin-resistance (*apr*) cassette containing *oriT*, which was amplified from pIJ773 using primers (TM8077/TM8078) with *lpdA*-specific extensions. The disrupted cosmid (pDI1) was confirmed by PCR analyses using the primers TM8079/TM8080 and TM5667/TM8080 and introduced into *E. coli* ET12567/pUZ8002 for conjugation into *S. venezuelae*. The *lpdA* mutants generated by double crossover were identified by their apramycin-resistant, kanamycin-sensitive phenotype. A representative null mutant was designated TMS0210 after confirmation by PCR using test primers TM8079/TM8080 and TM5667/TM8080.

For complementation, the native promoter and wild type coding sequence of *lpdA* (*vnz_09030*) were amplified using the primers TM8213/TM8243 and cloned into HindIII/Eco81I-cut pIJ10770 to generate pDI2 (pIJ10770-*lpdA*).^61^ The complemented plasmid was confirmed by PCR analyses using the primers TM5162/TM4359 and introduced into the *E. coli* ET12567/pUZ8002 for conjugation into TMS0210.

### Heterologous expression and purification of LpdA

The *lpdA* coding sequence (*vnz_09030*) was PCR-amplified using primers TM8196/ TM8197 and TM8198/TM8199 from genomic DNA from *S. venezuelae* NRRL B-65442 and cloned into pET28b(+) vector by Gibson assembly according to the standard protocol.^75^ Here, *lpdA* was ligated into the multiple cloning site of a modified pET28b(+) expression vector under the control of an IPTG-inducible T7 promoter, thereby fusing the gene to a sequence coding for once with C-terminal and once with an N-terminal 6×His fusion (pDI4 and pDI3, respectively). The complemented plasmid was confirmed by PCR analyses using the primers (TM6825/TM8203) and sequencing by Eurofins Genomics (Ebersberg, Germany).

The pDI3 or pDI4 plasmids were introduced into *E. coli* BL21 (DE3) cells for recombinant gene expression. LB medium was inoculated with cells of an overnight culture to a final OD_600_ of 0.1 and incubated at 37 °C with agitation at 220 rpm until an OD_600_ of 1 was reached. Expression was subsequently induced by addition of 0.1 mM IPTG and the temperature was reduced to 28 °C. Cells were harvested after 21 h by centrifugation (4 °C, 8000 rpm, 20 min, storage at −20 °C) before being washed and resuspended in ice-cold 20 mM Tris-HCl buffer (pH 8.0) containing 1 mM PMSF, 1 mg⋅mL^-1^ lysozyme, and 5 µg⋅mL^-1^ DNaseI. After incubation for 90 min at 6 °C and sonication for 5× 60 s with 60 s of rest between each sonication round (Sartorius Labsonic M, Settings: Cycle=0.6, Amplitude=100%), cell debris was removed by centrifugation (4 °C, 17000 rpm, 20 min). To purify the soluble N- or C-terminally His-tagged LpdA, the supernatant was filtered through a 0.45-micron filter and loaded onto a 5 mL HisTrap HP column (GE HealthcareLife Sciences, Chicago, IL, USA) connected to an ÄKTA FPLC system (GE Healthcare Life Sciences, Chicago, IL, USA). After washing with wash buffer (50 mM Tris-HCl (pH 8.0), 300 mM NaCl, and 10 mM imidazole), the bound protein molecules were eluted by increasing the imidazole concentration using elution buffer (50 mM Tris-HCl (pH 8.0), 300 mM NaCl, and 500 mM imidazole). The fractions with the highest LpdA concentrations were pooled and purified using HiPrep Desalting columns (GE HealthcareLife Sciences, Chicago, IL, USA) using storage buffer (50 mM Tris-HCl (pH 8.0), and 300 mM NaCl). The N- or C-terminally His-tagged LpdA proteins were tested for activity and analyzed by SDS-PAGE. Protein concentration of cell lysates was determined using the bicinchoninic acid assay (Pierce™ BCA Protein Assay Kits, Thermo Fisher Scientific™, Waltham, USA). Proteins were stored at 4 °C or –80 °C after addition of glycerol to a final concentration of 18%.

### Evaluation of protein fluorescence

For quantification of fluorescence spectra and intensities, the N- or C-terminally His-tagged LpdA proteins were analyzed using the Synergy NEO 2 HTS plate reader (BioTek, Winooski, VT, USA) with monochromators for illumination. The proteins were diluted to a final concentration of 50 µM and 150 µL were pipetted into a 96 well plate (black, clear bottom; Greiner Bio-One, Frickenhausen, Germany) followed by overhead readings. 50 µM FAD and a denatured LpdA sample were used as a control for the recorded excitation and emission spectra. To determine the effects of cofactor FAD on protein fluorescence, FAD cofactor was added to the LpdA samples (final concentration 50 µM), and fluorescence of each sample was measured using standard mCherry (EX:579/EM:616), and eGFP (EX:479/EM:520) filter sets. To record emission spectra, LpdA was illuminated at a fixed excitation wavelength (480 nm), and emitted fluorescence was recorded from 500 nm to 670 nm in 5 nm increments. This process was repeated for excitation spectra in 5 nm increments from 300 nm to 490 nm, and emission was measured at a fixed wavelength (520 nm). Mean values and standard deviation were determined from at least biological triplicates.

### Determination of the specific activity of isolated LpdA

The method used to evaluate LpdA activity concerns the reaction, i.e., the NADH-dependent reduction of lipoamide acid.^31^ The activity was determined spectrophotometrically using Cary 60 UV-Vis (Agilent Technologies, Santa Clara, US) at 22 °C by measuring the decrease of absorbance at 340 nm with 0.3 mM lipoamide acid and 0.2 mM NADH in 50 mM sodium phosphate buffer (pH 7.4) as previously described.^24^ One unit of enzyme activity was defined as the amount of enzyme that converted 1 µmol NADH per min. Specific activities were expressed as unit⋅mg^-1^ of protein, using 6220 M^-1^⋅cm^-1^ of molar absorption coefficient.

### Protein alignment

Bacterial, archaeal, and eukaryotic proteomes were analyzed for the LPDs. Amino acid sequences with high similarity to LpdA (*vnz_09030*) from *S. venezuelae* NRRL B-65442 were obtained from the National Center for Biotechnology Information (NCBI) and aligned to the LpdA from *S. venezuelae* using CLC Main Workbench 7 (Qiagen, Hilden, Germany) (Figure S6). A total of 25 protein sequences were aligned to construct phylogenetic trees. These included 16 bacterial, four archaeal, and five eukaryotic sequences (Table 3). A maximum likelihood tree using the Kimura Protein model with 100 bootstraps was applied to 25 aligned proteins provided. The final visualization of the phylogenetic trees was performed using iTOL.^76^

**Table 3.** LPD-proteins (NCBI-ID) used in this study.

| NCBI-ID | Organisms | Protein |
| --- | --- | --- |
| **Actinobacteria** |  |  |
| CCA55129 | *Streptomyces venezuelae*<br>NRRL B-65442 | Dihydrolipoyl dehydrogenase |
| CAB51264 | *Streptomyces coelicolor* A3(2) | Dihydrolipoyl dehydrogenase |
| WP_191868810 | *Streptomyces virginiae* | Dihydrolipoyl dehydrogenase |
| WP_156694758 | *Streptomyces ficellus* | Dihydrolipoyl dehydrogenase |
| WP_185023420 | *Actinomadura coerulea* | Dihydrolipoyl dehydrogenase |
| CNG84445 | *Mycobacterium tuberculosis* | Dihydrolipoyl dehydrogenase |
| **Proteobacteria** |  |  |
| NOJ79757 | *Myxococcus xanthus* | Dihydrolipoyl dehydrogenase |
| WP_006107278 | *Brucella abortus* | Dihydrolipoyl dehydrogenase |
| WP_002241465 | *Neisseria meningitidis* | Dihydrolipoyl dehydrogenase |
| AAC73227.1 | *Escherichia coli* K-12 substr. MG1655 | Lipoamide dehydrogenase |
| WP_317066271 | *Hydrogenimonas thermophila* | NAD(P)/FAD-dependent<br>oxidoreductase |
| **Firmicutes** |  |  |
| MCG0585108 | *Lactiplantibacillus plantarum* | Dihydrolipoyl dehydrogenase |
| HDJ1234727 | *Staphylococcus aureus* | Dihydrolipoyl dehydrogenase |
| WP_078175446 | *Bacillus mycoides* | Dihydrolipoyl dehydrogenase |
| WP_003230317 | *Bacillus* sp. | Dihydrolipoyl dehydrogenase |
| WP_191597703 | *Clostridium botulinum* | Dihydrolipoyl dehydrogenase |
| **Archaea** |  |  |
| WP_011250432 | *Thermococcus kodakarensis* | FAD-dependent<br>oxidoreductase |
| WP_008419176 | *Halalkalicoccus jeotgali* | Dihydrolipoyl dehydrogenase |
| WP_011012679 | *Pyrococcus furiosus* | Dihydrolipoyl dehydrogenase |
| WP_048043152 | *Methanosarcina mazei* | Dihydrolipoyl dehydrogenase |
| **Eukaryota** |  |  |
| AJU39818 | *Saccharomyces cerevisiae* YJM1252 | Dihydrolipoyl dehydrogenase |
| GAQ39161 | *Aspergillus niger* | Dihydrolipoyl dehydrogenase |
| NP_566570 | *Arabidopsis thaliana* | Lipoamide dehydrogenase 2 |
| EAA30299 | *Neurospora crassa* OR74A | Dihydrolipoyl dehydrogenase |
| 6HG8_A | *Homo sapiens* | Dihydrolipoyl dehydrogenase |

### Tracking the FOCI during the vegetative growth of hyphae

Fluorescent FOCI tracking was performed on microscopic images of single-cell microfluidic cultures using TrackMate software.^77^ The individual signals were detected by a DoG detector with an estimated object diameter of 0.6 microns and a sub-pixel localization function. The Advanced Kalman Tracker was used for frame-to-frame matching (initial search radius of 3.5 microns, search radius of 3.0, maximum frame gap of 2 frames, and feature penalty for quality (1.0)).

### Quantification analysis of microfluidic single-cell cultivation experiments

To further characterize the active growth and dynamics of the foci, the images of single-cell microfluidic cultures were analyzed at each time point (up to 600 min) using the open-source platform Fiji.^64^ To determine the increase in biomass as area [µm^2^], as well as the area [µm^2^] of the foci and their intensity, the growing hyphae were identified using the automatic area selection of Fiji (threshold at 10%). The “Analyze Particles” function was used to identify the region of interest (ROI) (threshold at 0.032 µm^2^) and subsequent analysis using the “Measure” function (Supporting Information S6). The number of the foci at each time point, as well as the sum, mean of the area [µm^2^], and intensity at each time point, were calculated and plotted for all recorded values. The specific increase of biomass as growth rate [µm^2^/min] and the average distance of FOCI [n(FOCI)/min] were calculated from the log growth phase according to the following formula:

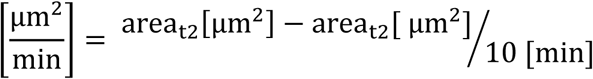

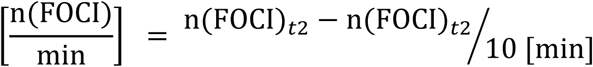

The values were then normalized to the number of actively growing tips of hyphae to determine the specific values per tip [(µm^2^/minute)/tip] and [(n(FOCI)/min)/tip].

### Agent-based model for the growth and branching behavior of *Streptomyces* spp. mediated by LpdA-containing foci

To explore the implications of localized metabolism in *Streptomyces*, we developed an agent-based model that simulates hyphal growth and branching influenced by LpdA-containing foci. By incorporating cellular components including discrete foci as sites of localized ATP production, polarisome for tip extension, and continuous fields representing ATP and macromolecule gradients, the model successfully reproduces experimentally observed growth patterns and foci distributions.

The multicellular cell body is represented by a collection of non-overlapping circular segments and serve as a domain where the reaction-diffusion processes happen. Growth of the filament happens by addition of the new segments at the tips of the filament or at the branching points. Fluorescent foci are represented by circular particles that perform Brownian diffusion. Foci grow from a certain minimum to a fixed maximum size. Foci are also affected by the drift due to cytoplasmic streaming during the filament tip growth. Foci serve as a source of energy in the form of ATP molecules. ATP molecules are thus produced in the segments containing foci with a rate proportional to the foci size. ATP is converted to produce macromolecules. It is assumed that ATP consumption rate is higher towards the growing tip. Macromolecules in the presence of the polarisome, are consumed to make new cellular material. Once enough of the cellular material is accumulated, the new filament segment is added. Conversion of the macromolecules to the new cellular segments is accomplished via and proportional to the size of the polarisome, which are modeled to be present in every tip and at branching points. Like foci, polarisomes grow and at a certain size can randomly split leaving a fraction behind that will later initiate filament branching point. Concentration of ATP is modeled by a reaction diffusion equation, where ATP is produced by foci and converted into macromolecules and consumed at basal rate for housekeeping needs. Marcomolecules are being produced in segments as a process of ATP conversion, are consumed to build new segments, and are degraded at a certain rate. Diffusion equations for ATP and macromolecules are discretized down to the individual segment level. Initial values of many parameters of the model are based on the literature values and tuned to match the experimental observations on the filament growth and density of the foci per filament length.

Our simulations yield growth rates (0.46 h^-1^) and foci densities (0.70 foci/µm) that closely match experimental measurements (0.38-0.47 h^-1^) and (0.66-0.72 foci/µm), respectively. The simulation results show heterogeneity in ATP and macromolecule concentrations throughout the hyphae. This heterogeneity arises from the interplay between localized ATP production at foci sites, random motion of foci, and elevated consumption near growing tips, where energy-intensive processes like cell wall synthesis occur. The model’s ability to reproduce both normal growth patterns and phenotypic variations like hyperbranching by parameter adjustments highlights it usefulness for studying the impact of energy metabolism and macromolecule consumption on the multicellular organization of *Streptomyces* species. For a detailed description of the model and its parameters please see Supporting Information S8.

