## supplementary figurs for "Spatiotemporal Dynamics of the Dihydrolipoyl Dehydrogenase LpdA Determine the Intrinsic Autofluorescence in Filamentous Actinobacteria"

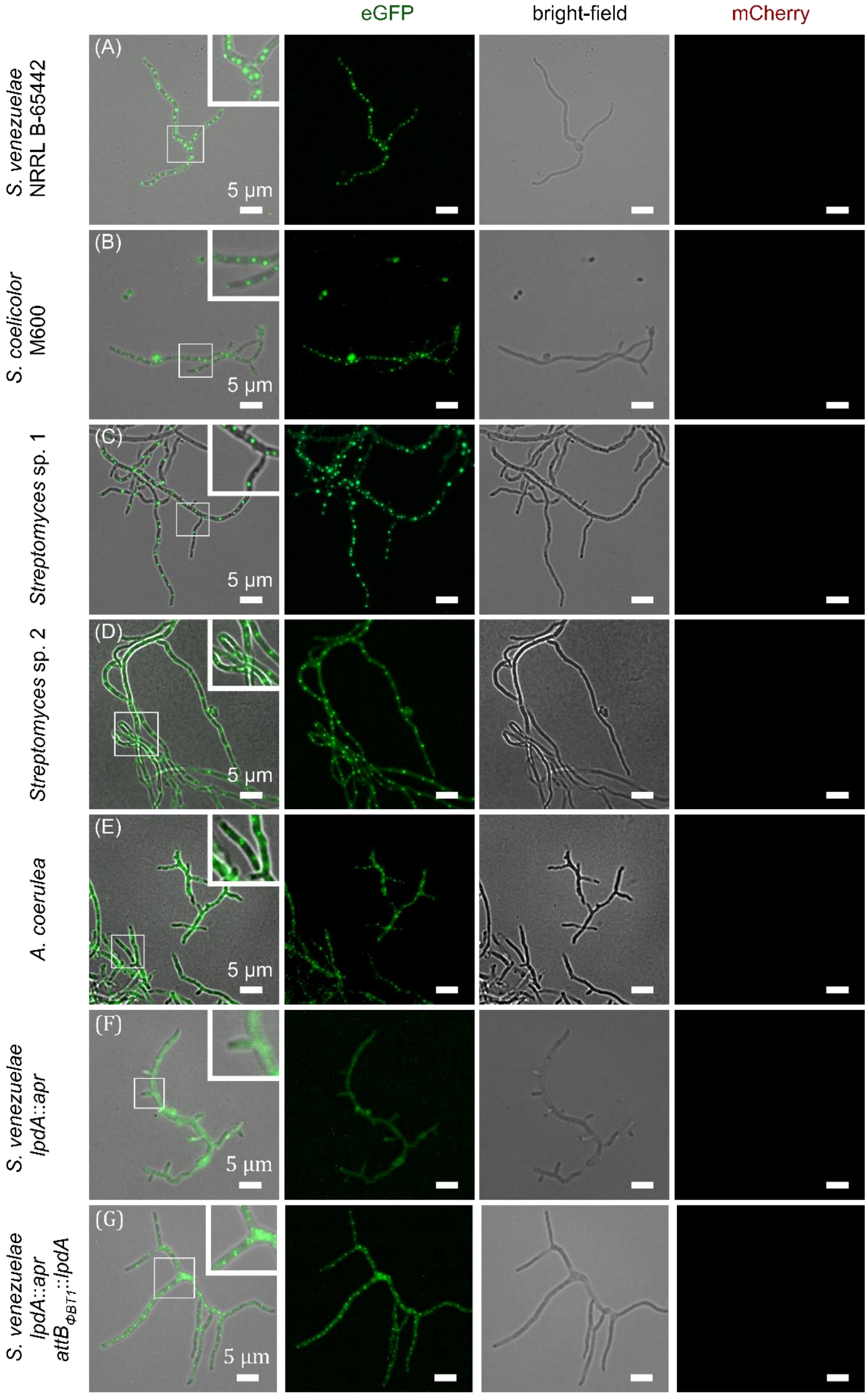


**Figure S1. Fluorescence microscopy of *S. venezuelae* NRRL B-65442, *S. coelicolor* M600, *A. coerulea,* and two *Streptomyces* sp. (1 and 2) isolated from the environment.** Fluorescence microscopy of the hyphae was conducted using an Axio Observer 7 inverted microscope (Carl Zeiss, Jena, Germany) equipped with the standard eGFP ($\lambda_{EX}$ = 488 nm; $\lambda_{EM}$ = 509 nm) and mCherry ($\lambda_{EX}$ = 587 nm; $\lambda_{EM}$ = 610 nm; exposure time of 1.5 s) filter sets.

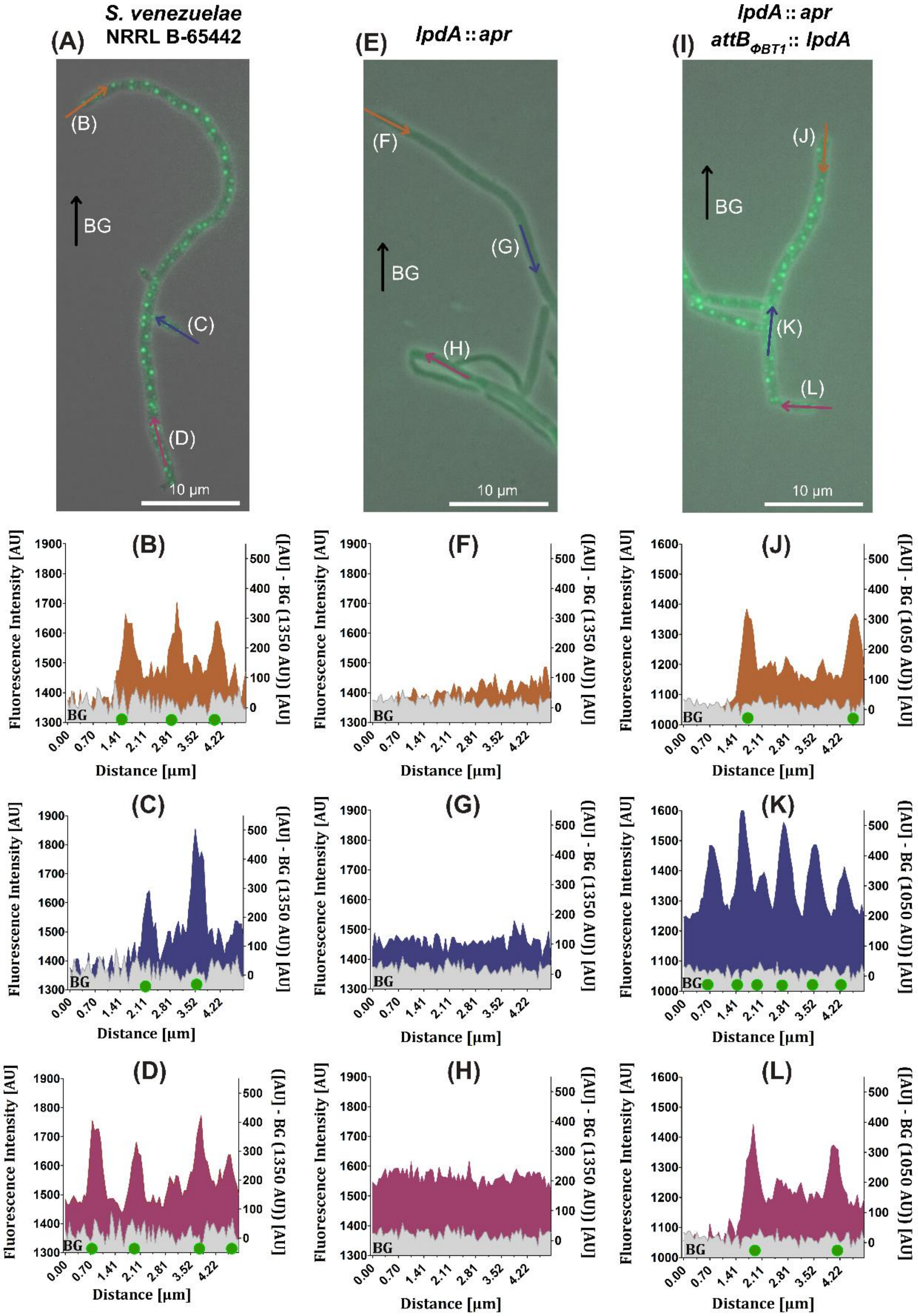


**Figure S2. Fluorescence intensity profiles for *S. venezuelae* NRRL B-65442 (WT), null mutant (*lpdA*::*apr*), and complemented null mutant (*lpdA*::*apr attB*$_{\Phi BT1}$::*lpdA*).** Fluorescence microscopy of the hyphae was conducted using an Axio Observer 7 inverted microscope (Carl Zeiss, Jena, Germany) equipped with the standard eGFP ($\lambda_{EX}$ = 488 nm; $\lambda_{EM}$ = 509 nm) and mCherry ($\lambda_{EX}$ = 587 nm; $\lambda_{EM}$ = 610 nm; exposure time of 1.5 s) filter sets. Original microscopy images were utilized for analysis. Pixel intensity was quantified at three positions along the hyphae (colored arrows). Background fluorescence of the agarose pads was quantified as a control (black arrow) and depicted in gray on the graphs. Green circles correspond to the position of the detected hyphal foci in the analyzed areas.

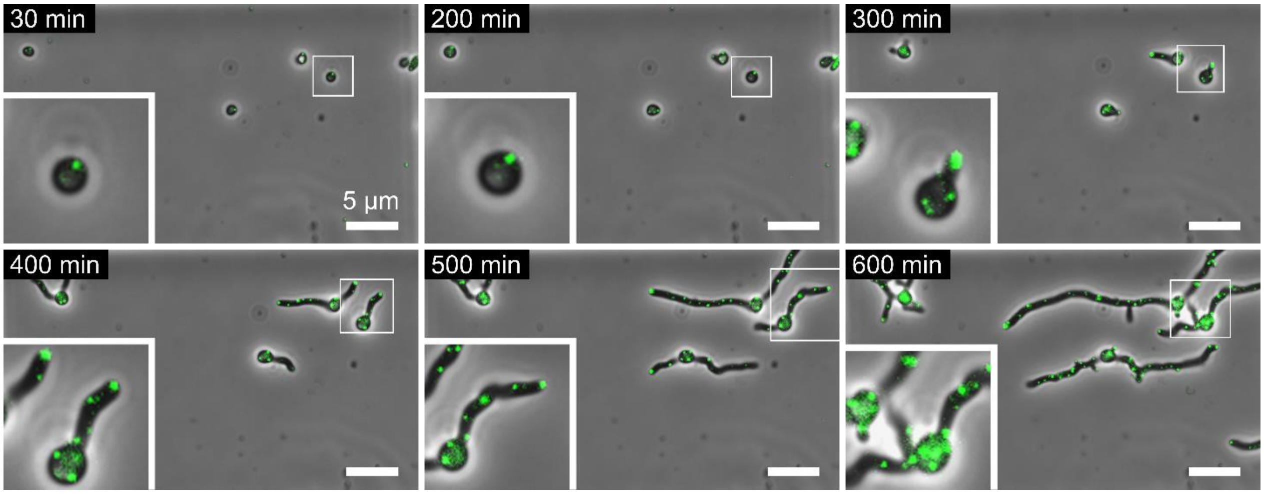


**Figure S3. Microfluidic single-cell cultivation (MSCC) under continuous time-lapse imaging for *S. coelicolor* M600.** The strain was injected into the MSCC system and cultivated at 28 °C for 12.5 h with images being taken in 10 min intervals. Fluorescence microscopy of the hyphae was conducted using an Axio Observer 7 inverted microscope (Carl Zeiss, Jena, Germany) equipped with the standard FITC_38 filter set ($\lambda_{EX}$ = 495 nm; $\lambda_{EM}$ = 519 nm). Illumination times for phase contrast and fluorescence microscopy images were 40 ms and 700 ms, respectively. Excitation light transmission was set to 25%. Red arrows indicate branching points. For the corresponding movies, see Supporting Information Movies.

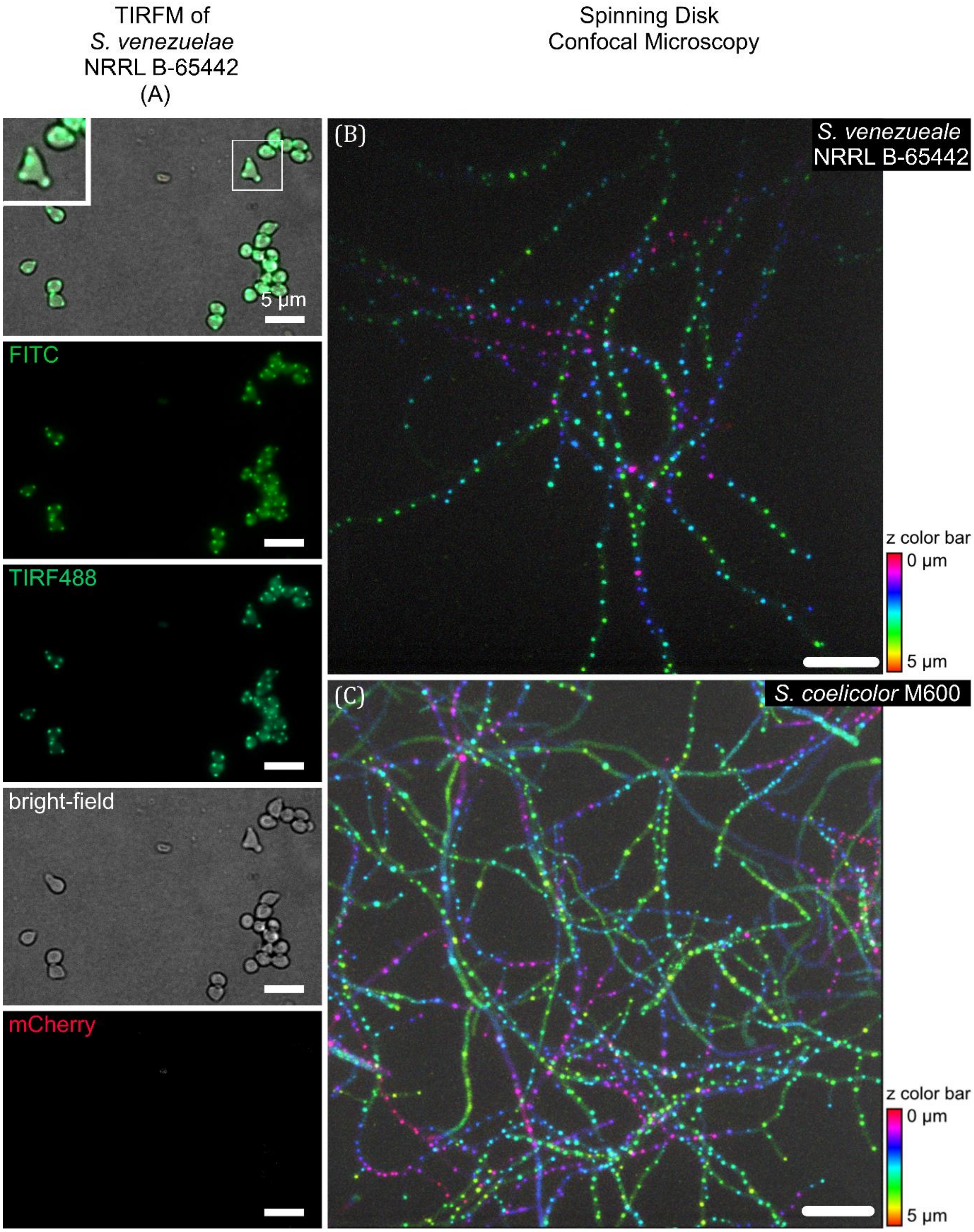


**Figure S4. (A) TIRF microscopy on germinated *S. venezuelae* NRRL B-65442 spores.** Spores were incubated at 28 °C for four hours before being utilized for TIRF microscopy with a DeltaVision system. Images were captured with an Olympus 1.45 NA 100× objective lens, the standard FITC ($\lambda_{EX}$ = 495 nm; $\lambda_{EM}$ = 519 nm) and mCherry ($\lambda_{EX}$ = 587 nm; $\lambda_{EM}$ = 610 nm; exposure time of 1.5 s) filter sets, and an argon ion laser source of wavelength 488 nm (TIRF488). **Spinning Disk Confocal Microscopy of *S. venezuaelae* NRRL B-65442 (B) and *S. coelicolor* M600 (C).** For cellular fluorescence imaging, a 488 nm laser with 50 mW (measured before the objective) was used for excitation under HILO conditions, with an exposure time of 100 ms per frame, and a total of 100 framers were recorded.

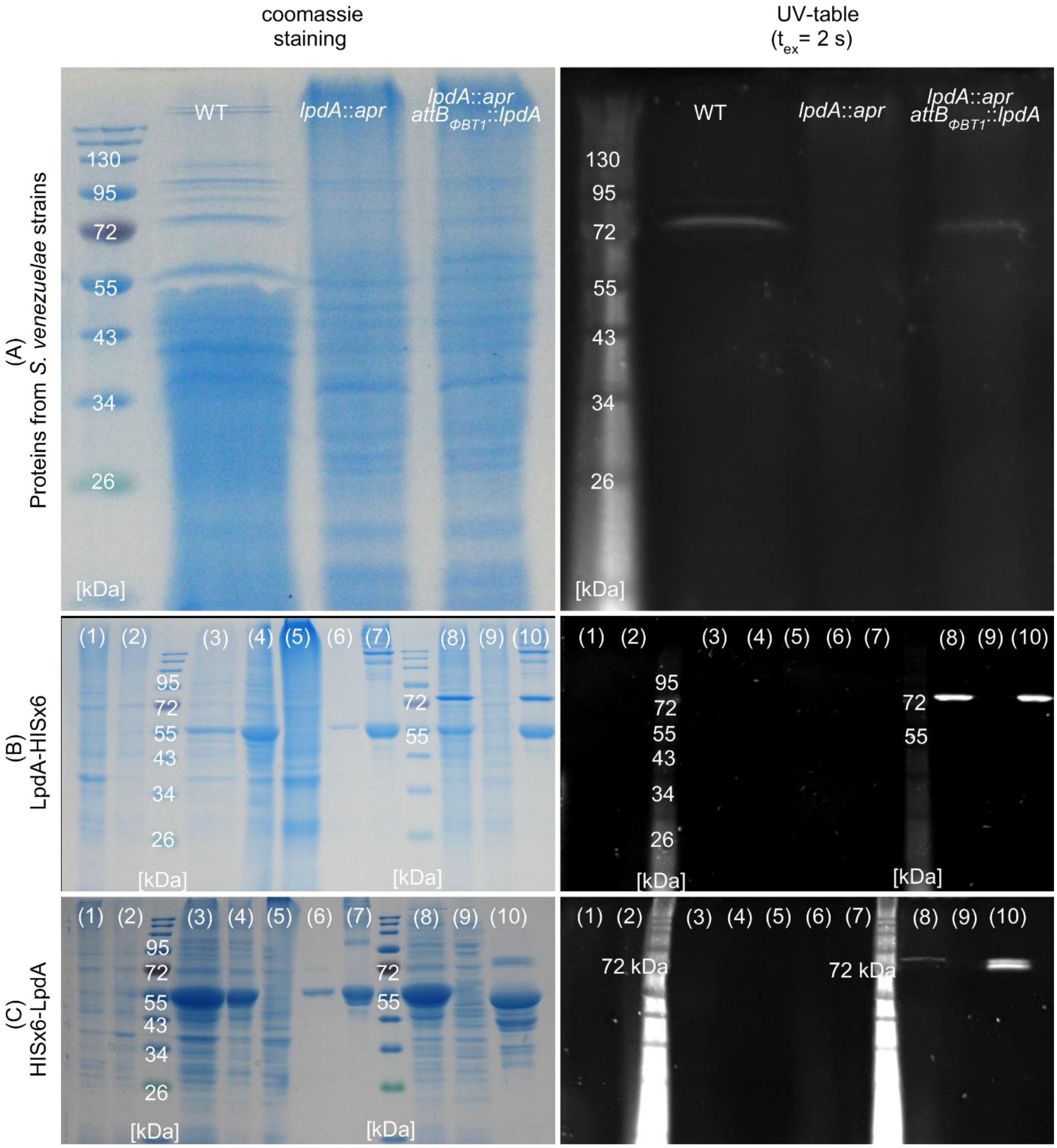


**Figure S5. SDS-PAGE of proteins from *S. venezuelae* and *lpdA* mutants (A) and *E. coli* BL21 expressed C- and N-terminally 6×His-tagged LpdA (B and C, respectively). (A)** Proteins were isolated from *S. venzuelae* after cultivation at 28 °C for 20 h. 15 µg of the isolated proteins per lane were used for SDS-PAGE gel. **(B) and (C)** Lanes 1 and 2: Denatured proteins from *E. coli* BL21 before induction of LpdA-6×His expression via the addition of IPTG (lane 1: total cell lysate; lane 2: supernatant). Lanes 3 and 4: total cell lysate and supernatant fractions, respectively, of the BL21-harboring pET28b-LpdA-6×His plasmid after IPTG-induction. Lanes 5 and 9: negative controls of denatured (5) and native (9) BL21 proteins. Lanes 6 and 8: LpdA-6×His after nickel affinity chromatography (after denaturation (6) and in native form (8)). Lanes 7 and 10: Purified LpdA-6×His after denaturation (7) and in native form (10). Arrows denote monomers (red) and dimers (green) of LpdA. Thermo PageRuler™ prestained protein ladder was used as a marker (M). Protein fluorescence was detected using a UV transilluminator. For colorimetric visualization, staining was performed with Coomassie brilliant blue R-250.

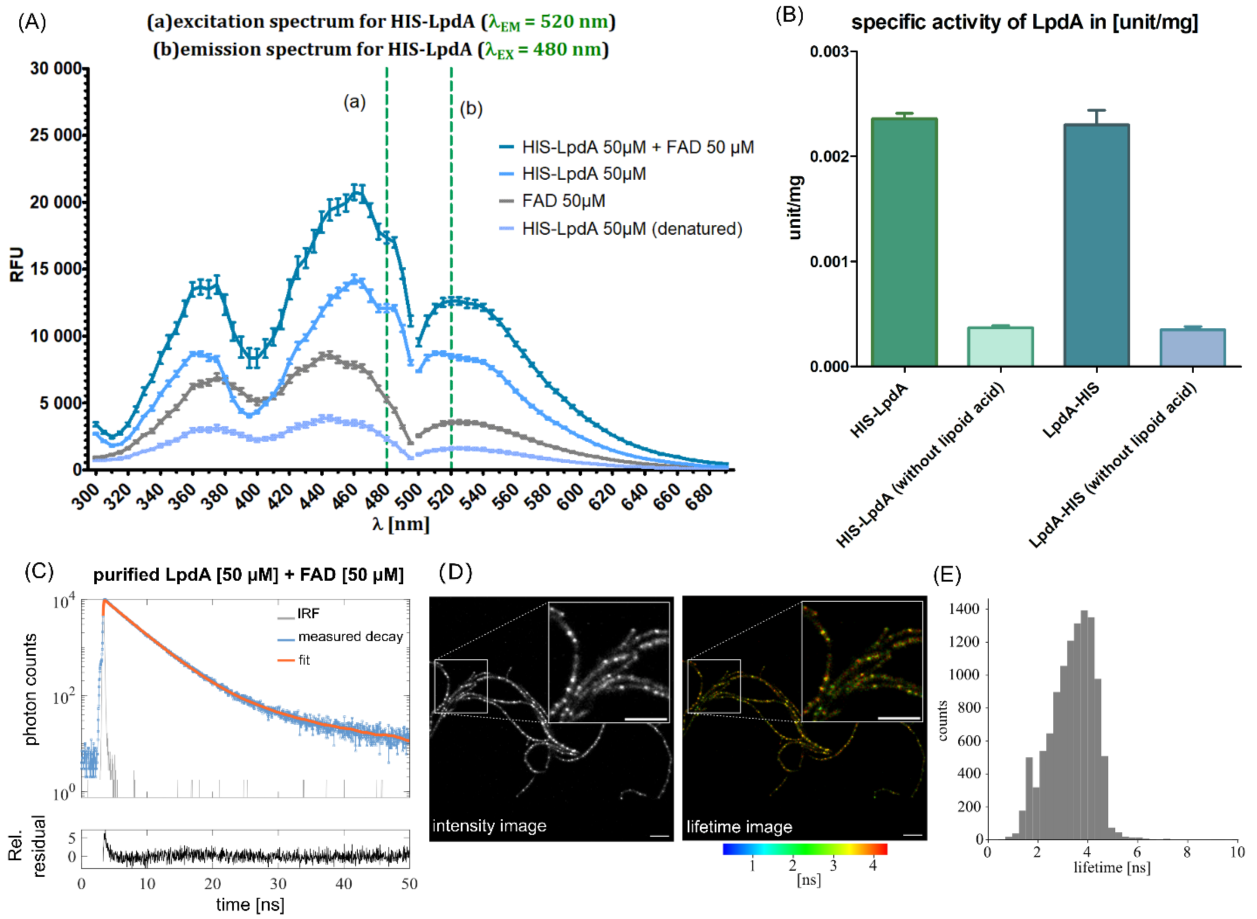


**Figure S6. Specific activity of purified LpdA (A) and fluorescence spectra of lipoamide dehydrogenase (6×His-LpdA) (B). (A)** Activity was determined spectrophotometrically using Cary 60 UV-Vis (Agilent Technologies, Santa Clara, US) at 22 °C by measuring the decrease of absorbance at 340 nm. One unit of enzyme activity was defined as the amount of enzyme that converted 1 µmol NADH per min. The reaction "without lipoid acid" was used as a negative control. **(B)** The excitation spectrum (a) was measured at 520 nm emission and the emission spectrum (b) was obtained at 480 nm excitation. Fluorescence spectra are shown for purified 50 µM 6×His-LpdA (blue), 50 µM FAD (gray), 50 µM denatured 6×His-LpdA (light blue), and 50 µM 6×His-LpdA after incubation with 50 µM FAD (dark blue). **The fluorescence lifetime decay of purified LpdA-6×His was obtained in a TCSPC spectrometer (C)**. The detected photon counts are shown in blue on a logarithmic scale. The decay was fitted with a biexponential fit (orange line). The instrument response function (IRF) is represented in grey. **Fluorescence lifetime imaging microscopy of cellular autofluorescence in living *S. venezuelae* (D) and (E).** Fluorescence intensity and lifetime images with a zoomed in view in the top right, scale bar: 5 µm. Lifetime range is represented from 0 ns (blue) to 4.3 ns (red). Panel (E) shows the fluorescence lifetime histogram corresponding to the field of view with a mean of 3.6 ± 0.1 ns.

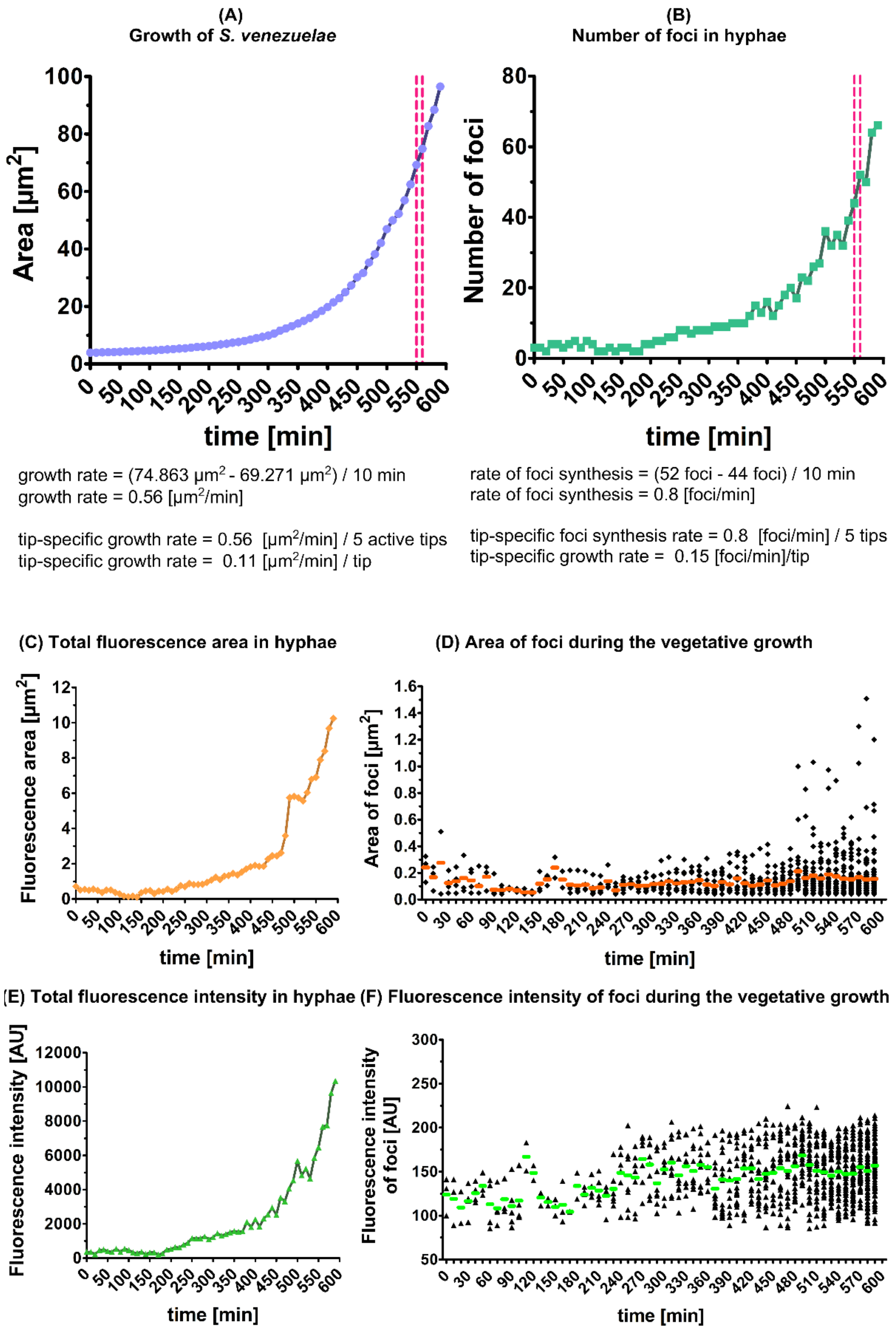


*Figure S 7.* **The quantitative analysis of the MSCC for *S. venezuelae* NRRL B-65442**. Data demonstrated a direct correlation between the active growth rate of the hyphae (A), the number of detected foci (B), the total fluorescence intensity (C), and the total fluorescence area (E). Furthermore, the area (D) and intensity (F) of the individual foci were displayed as a dot plot. The utilization of colored lines (green for fluorescence intensity and orage for area of foci) in the representation signifies the calculation of the mean values at specific temporal points. The red dashed lines indicate the period (550-560 min) used to calculate the growth rate and the rate of foci synthesis.

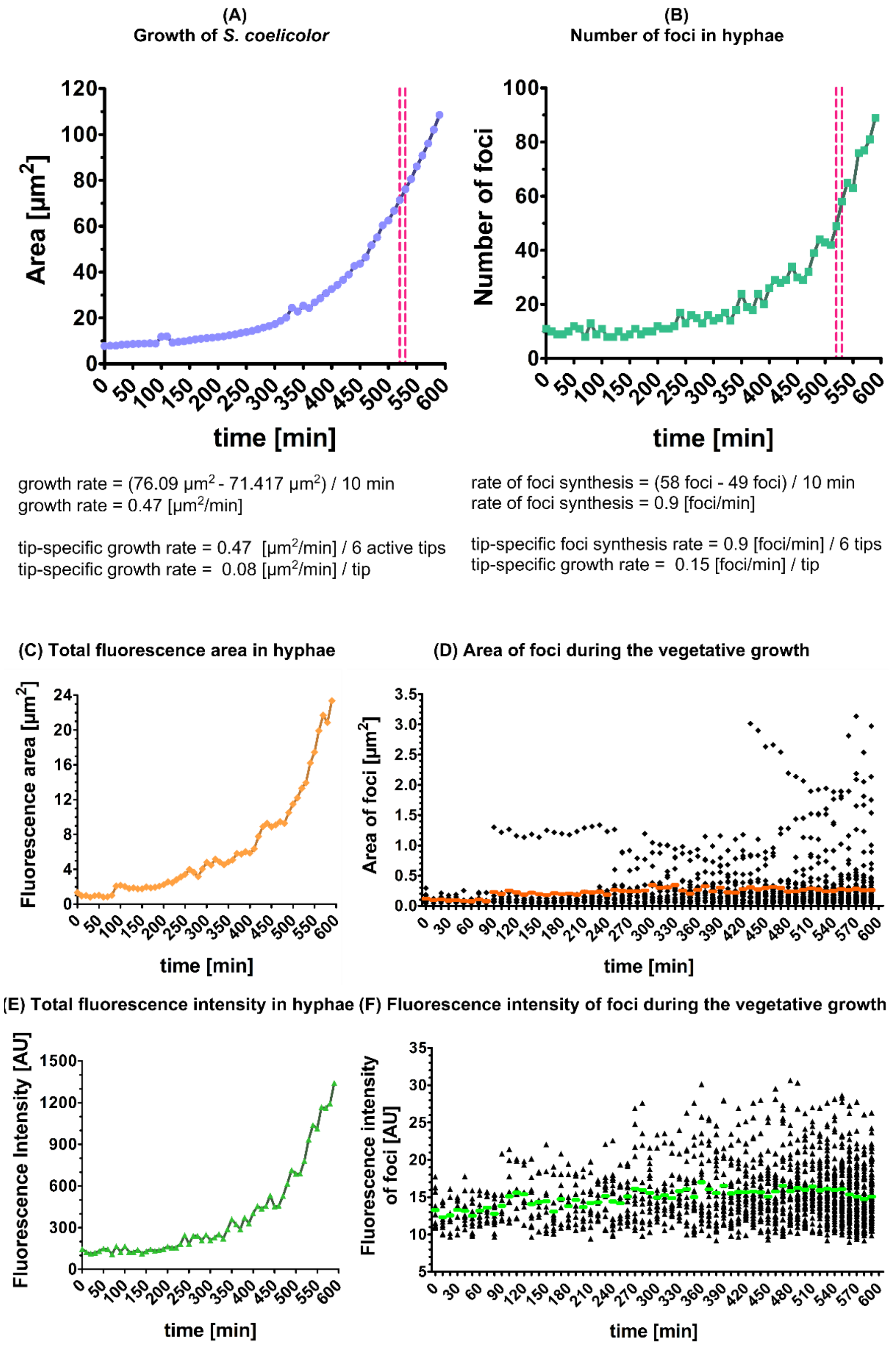


*Figure S 8* **The quantitative analysis of the MSCC for *S. coelicolor* M600**. Data demonstrated a direct correlation between the active growth rate of the hyphae (A), the number of detected foci (B), the total fluorescence intensity (C), and the total fluorescence area (E). Furthermore, the area (D) and intensity (F) of the individual foci were displayed as a dot plot. The utilization of colored lines (green for fluorescence intensity and orage for area of foci) in the representation signifies the calculation of the mean values at specific temporal points. The red dashed lines indicate the period (520-530 min) used to calculate the growth rate and the rate of foci synthesis.

***Table S1*** *Parameters obtained from FLIM analysis of cellular auto-fluorescence in living Streptomyces venezuelae. The table includes values for the three lifetime components with their relative amplitude and intensity as well as the mean intensity and mean amplitude weighted fluorescence lifetime.*

| **Parameter** | |
|---|---|
| **$\tau_1$ [ns]** | 0.3±0.1 |
| **$\tau_2$ [ns]** | 1.5±0.3 |
| **$\tau_3$ [ns]** | 4.7±0.4 |
| **intensity 1 [kCnts]** | 22.6±7 |
| **intensity 2 [kCnts]** | 123.4±25 |
| **intensity 3 [kCnts]** | 319.8±26 |
| **<$\tau$> int [ns]** | 3.6±0.1 |
| **χ2** | 1.040 |

***Table S2*** *Parameters obtained from TCSPC analysis of the fluorescence lifetime of purified* LpdA-6×His*. The parameters were extracted from a two component reconvolution fitting of the decay curves and include the two fluorescence lifetimes $\tau$1, $\tau$2, the corresponding fraction f1 of the first component, and the mean amplitude and mean intensity weighted lifetimes.*

| **Parameter** | **LpdA-6×His + FAD** |
|---|---|
| **$\tau$ 1 [ns]** | 3.7±0.13 |
| **$\tau$ 2 [ns]** | 13.6±1.8 |
| **f1** | 0.93±0.2 |
| **<$\tau$> amp [ns]** | 4.47 |
| **<$\tau$> int [ns]** | 5.98 |

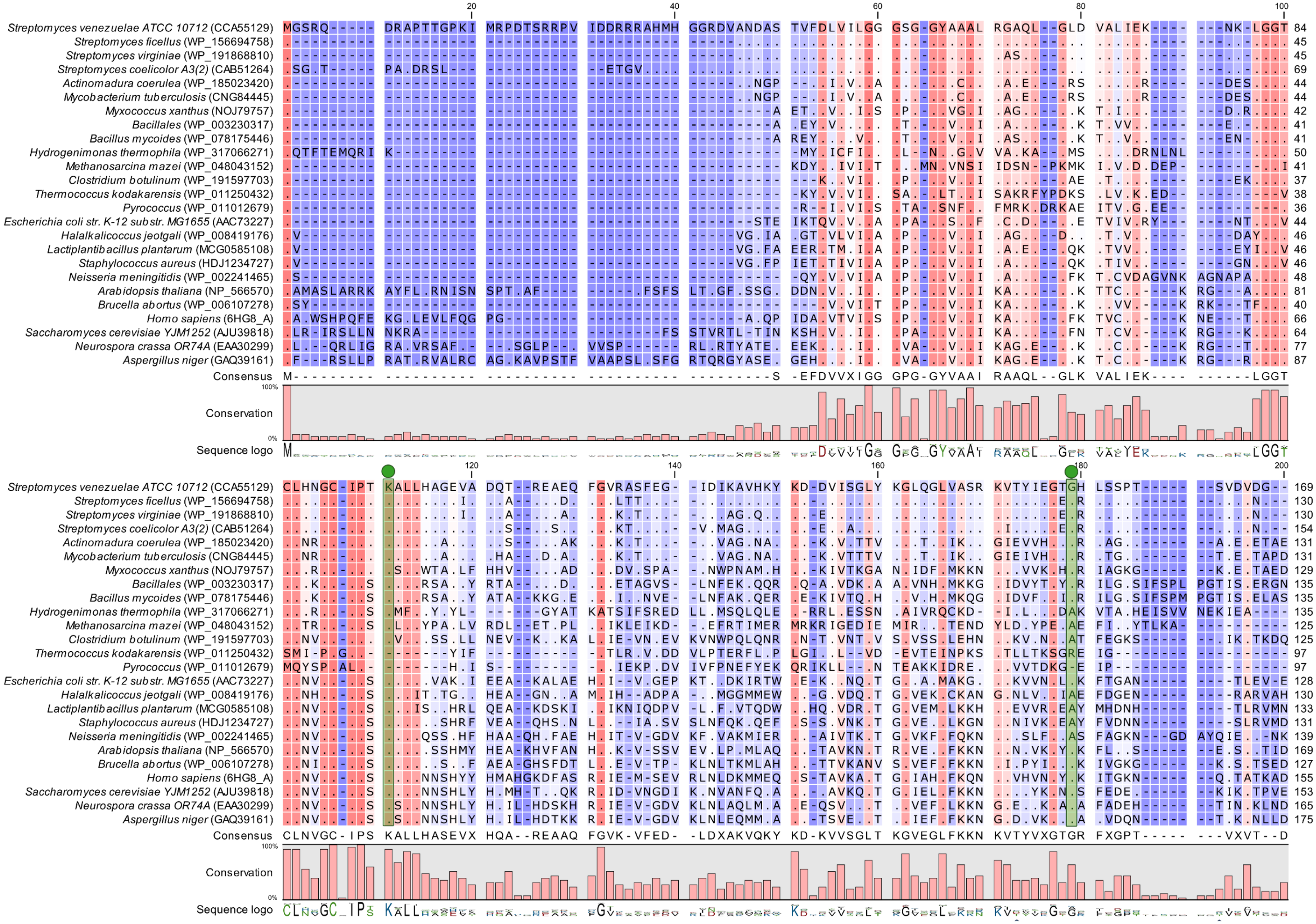

20
40
60
80
100
Streptomyces venezuelae ATCC 10712 (CCA55129)
Streptomyces ficellus (WP_156694758)
Streptomyces virginiae (WP_191868810)
Streptomyces coelicolor A3(2) (CAB51264)
Actinomadura coerulea (WP_185023420)
Mycobacterium tuberculosis (CNG84445)
Myxococcus xanthus (NOJ79757)
Bacillales (WP_003230317)
Bacillus mycoides (WP_078175446)
Hydrogenimonas thermophila (WP_317066271)
Methanosarcina mazei (WP_048043152)
Clostridium botulinum (WP_191597703)
Thermococcus kodakarensis (WP_011250432)
Pyrococcus (WP_011012679)
Escherichia coli str. K-12 substr. MG1655 (AAC73227)
Halalkalicoccus jeotgali (WP_008419176)
Lactiplantibacillus plantarum (MCG0585108)
Staphylococcus aureus (HDJ1234727)
Neisseria meningitidis (WP_002241465)
Arabidopsis thaliana (NP_566570)
Brucella abortus (WP_006107278)
Homo sapiens (6HG8_A)
Saccharomyces cerevisiae YJM1252 (AJU39818)
Neurospora crassa OR74A (EAA30299)
Aspergillus niger (GAQ39161)
Consensus M--------- ---------- ---------- ---------- ---------S -EFDVVXIGG GPG-GYVAAI RAAQL--GLK VALIEK---- ------LGGT
Conservation
100%
0%
Sequence logo
120
140
160
180
200
Consensus CLNVGC-IPS KALLHASEVX HQA--REAAQ FGVK-VFED- -LDXAKVQKY KD-KVVSGLT KGVEGLFKKN KVTYVXGTGR FXGPT----- ---VXVT--D
Conservation
Sequence logo

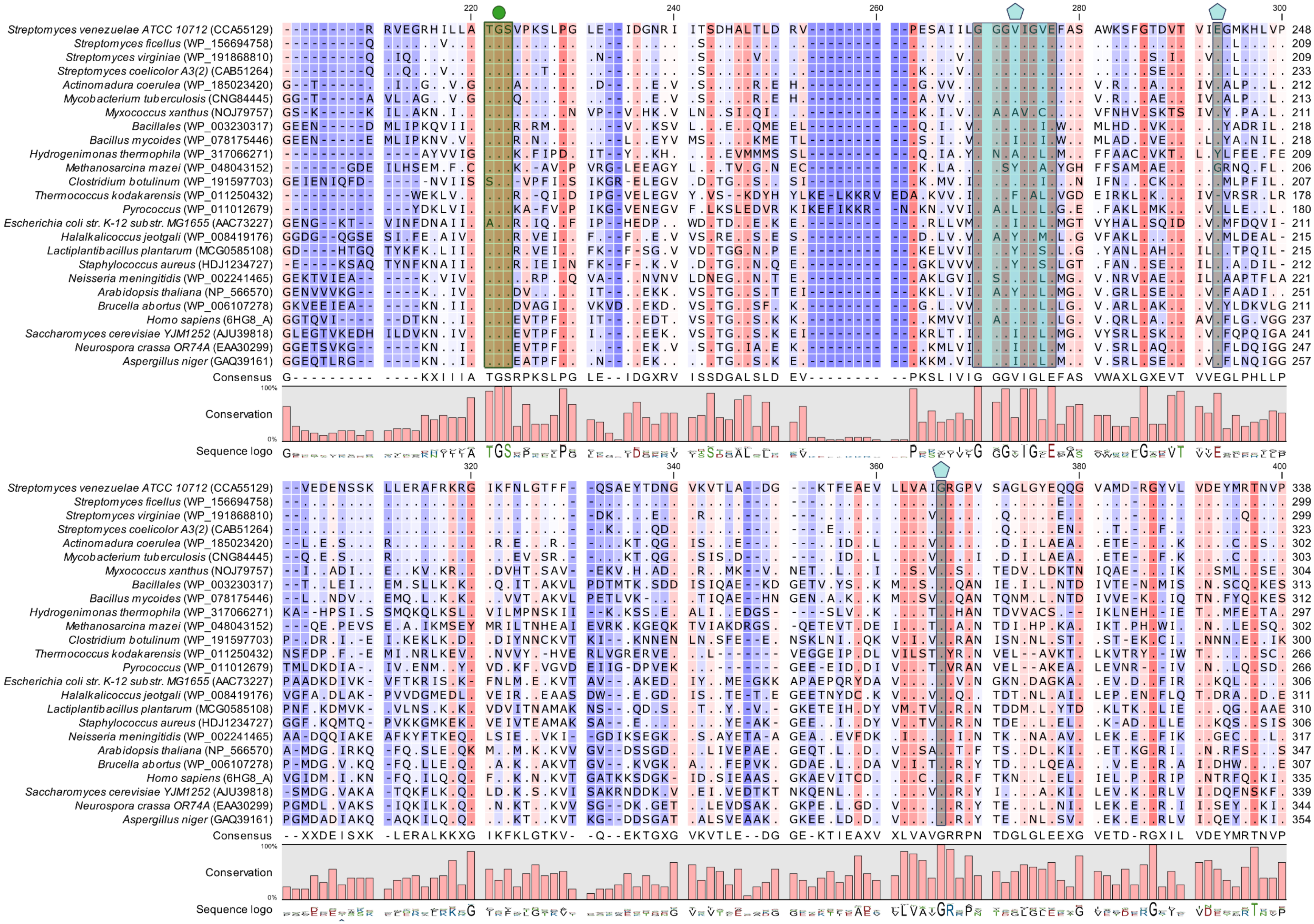

Streptomyces venezuelae ATCC 10712 (CCA55129)
Streptomyces ficellus (WP_156694758)
Streptomyces virginiae (WP_191868810)
Streptomyces coelicolor A3(2) (CAB51264)
Actinomadura coerulea (WP_185023420)
Mycobacterium tuberculosis (CNG84445)
Myxococcus xanthus (NOJ79757)
Bacillales (WP_003230317)
Bacillus mycoides (WP_078175446)
Hydrogenimonas thermophila (WP_317066271)
Methanosarcina mazei (WP_048043152)
Clostridium botulinum (WP_191597703)
Thermococcus kodakarensis (WP_011250432)
Pyrococcus (WP_011012679)
Escherichia coli str. K-12 substr. MG1655 (AAC73227)
Halalkalicoccus jeotgali (WP_008419176)
Lactiplantibacillus plantarum (MCG0585108)
Staphylococcus aureus (HDJ1234727)
Neisseria meningitidis (WP_002241465)
Arabidopsis thaliana (NP_566570)
Brucella abortus (WP_006107278)
Homo sapiens (6HG8_A)
Saccharomyces cerevisiae YJM1252 (AJU39818)
Neurospora crassa OR74A (EAA30299)
Aspergillus niger (GAQ39161)
Consensus
G- - - - - - - - - - - - KXIIIA TGSRPKSLPG LE- - IDGXRV ISSDGALSLD EV- - - - - - - - - - PKSLIVIG GGVIGLEFAS VWAXLGXEVT VVEGLPHLLP
Conservation
Sequence logo
Consensus
- - XXDEISXK -LERALKKXG IKFKLGTKV- -Q- -EKTGXG VKVTLE- -DG GE-KTIEAXV XLVAVGRRPN TDGLGLEEXG VETD-RGXIL VDEYMRTNVP
Conservation
Sequence logo

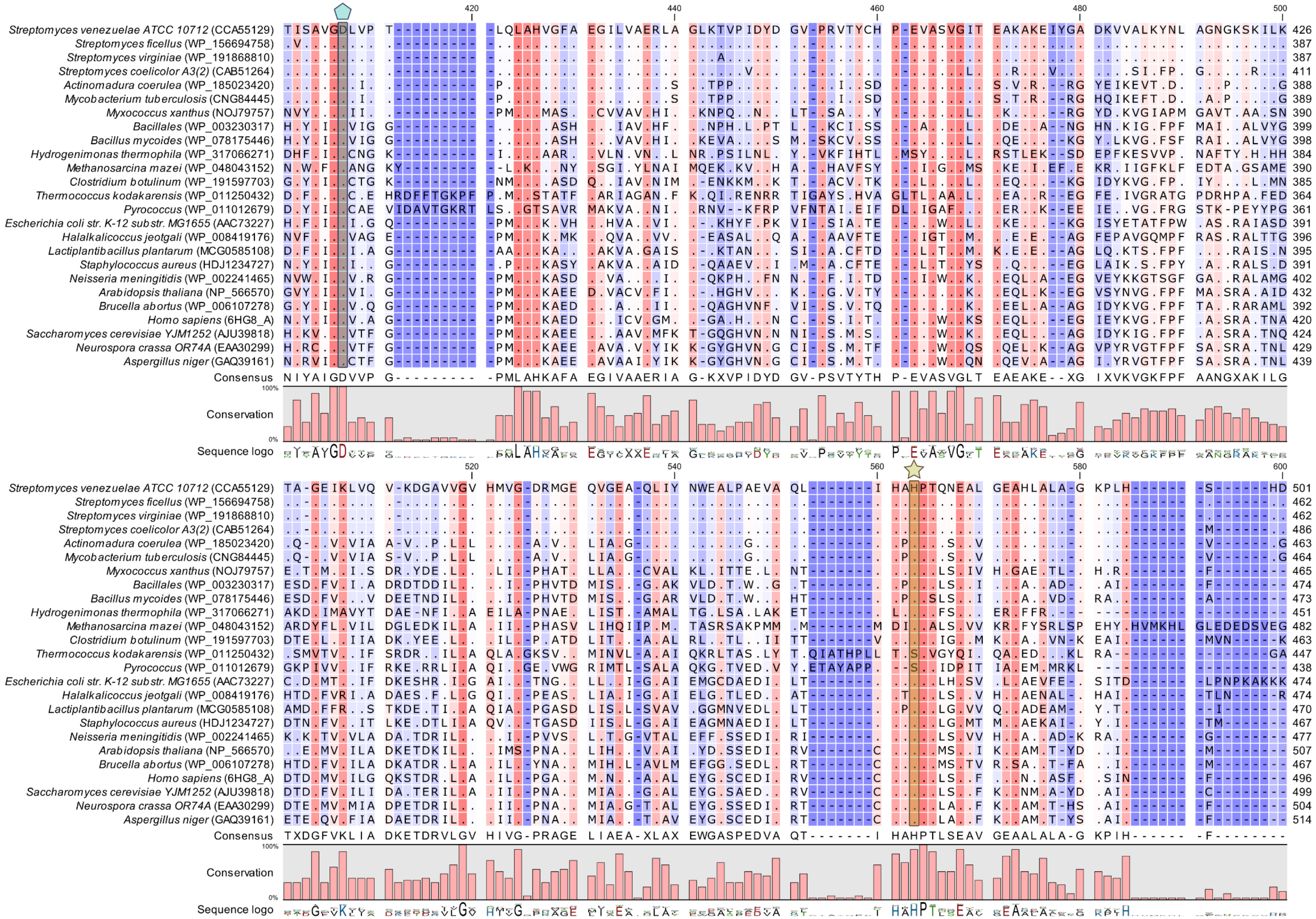


**Figure S8. Primary structure comparison between lipoamide dehydrogenase from *S. venezuelae* NRRL B-65442 (as ATCC 10712) and lipoamide dehydrogenases from various species.** Sequences were retrieved from GenBank and corresponding accession numbers are provided in Table 3. Identical residues are indicated by dots and sequence gaps by dashes. Residues are marked indicating FAD binding sites (green; circle), NADH binding sites (blue; pentagon) and proton acceptors (yellow; star). Color gradient indicates conservation (pink: 100%; dark blue: 0%).
