## Supplementary figures and images for "Spatiotemporal Dynamics of the Dihydrolipoyl Dehydrogenase LpdA Determine the Intrinsic Autofluorescence in Filamentous Actinobacteria"

### BF_S-coelicolor_60_frams.tif

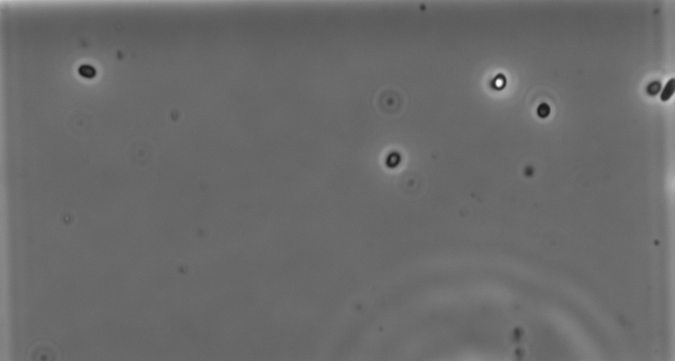

### BF_S. venezuelae.tif

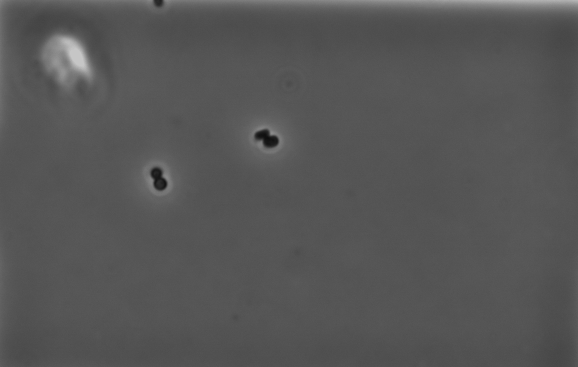

### FOCI_S. venezuelae.tif

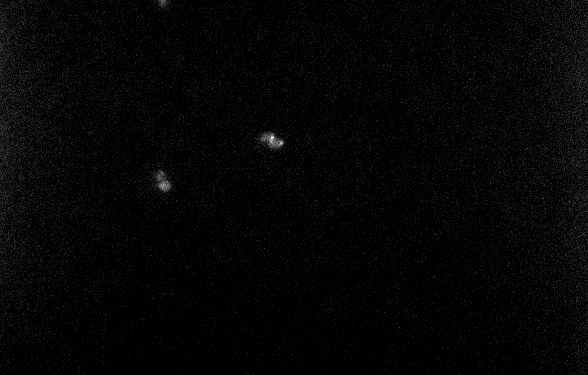
