## Supplementary material for "Spatiotemporal Dynamics of the Dihydrolipoyl Dehydrogenase LpdA Determine the Intrinsic Autofluorescence in Filamentous Actinobacteria": S8_Theory_for_agent_based_model

### Agent-based model for the growth and branching behavior of *Streptomyces* species mediated by LpdA-containing foci.

### 1. Introduction

Our experimental results demonstrate that green fluorescent foci within *Streptomyces* hyphae contain the lipoamide dehydrogenase A (LpdA) protein and that flavin adenine dinucleotides (FAD) when bound to LpdA, are the source of their green autofluorescence. Since LpdA plays a crucial role in the tricarboxylic acid (TCA) cycle as part of the multienzyme supercomplex PDH-ODH, these foci are hypothesized to be the sites of elevated metabolic activity.[1]

We propose that the green fluorescent foci could represent discrete intracellular locations where central energy metabolism pathways, such as the TCA cycle, are highly active, leading to localized production of metabolites including electron carriers NADH and $FADH_2$.[2] Given that the foci are closely associated with the cell membrane (at least at the tips and branching points of hyphae), we further hypothesized that the presence of foci could result in localized ATP synthesis through oxidative phosphorylation on the cell membrane. Similarly, the localization of foci to the growing tip of the hyphae suggests their potential role in ATP-dependent activities such as cell wall synthesis, hyphal extension, and branching.

To explore the possible repercussions of localized ATP production on *Streptomyces* morphology and multicellular organization, we have developed a phenomenological, agent-based, computational model describing hyphal growth and branching. This model considers key cellular components and their interactions, in particular focusing on the localization and movement of LpdA-containing foci, ATP gradients, macromolecules' gradients, and polarisome (or TIPOC (TIP Organizing Center)) splitting.

The model is built upon several key assumptions:

- **Localized metabolic activity:** Metabolic enzymes involved in the TCA cycle are not homogeneously distributed throughout the whole cytoplasm, but instead form protein complexes at specific discrete locations (foci) where metabolic activity is concentrated.
- **Fast NADH and $FADH_2$ consumption:** Electron carriers NADH and $FADH_2$, presumably produced at the foci sites, are rapidly utilized by the nearby electron transport chain. This allows us to assume that ATP is also produced in a localized manner in the close vicinity of foci.

- **Non-uniform ATP Consumption:** ATP is consumed at a higher rate in regions close to the growing tips, where the most energy-intensive processes such as protein synthesis and cell envelope synthesis occur.[3–5]

### 2. Overview of the Model

The components of the model can be separated into two main categories:

- **Cellular components modeled by autonomous agents:**
  These components are discrete entities that interact with each other:
  - **Cell segments (compartments):** Together they form a chain that represents the hyphae.
  - **Polarisomes:** Locations where new cell envelope material is added, leading to hyphal elongation or branching. They are always present at the tips of the hyphae and at branching points.
  - **LpdA-containing foci:** Modeled as Brownian particles where protein complexes involved in the TCA cycle are localized.
- **Cellular concentrations modeled by continuous fields:**
  These components are represented by continuous fields that evolve over time according to reaction-diffusion equations with discretization at the level of individual cell segments:
  - **ATP:** Required for synthesis of macromolecules and housekeeping processes.
  - **Macromolecules:** Required for tip elongation and branching.

Below we describe in more detail how each of the categories are implemented in the model. For simplicity, we consider the implementation of the model in a planar (two-dimensional) geometry.

### 3. Cellular components modeled by agents

#### 3.1. Cell segments

The hyphae are represented as a chain of interconnected cell segments (compartments) (Figure 1). Most of the segments are connected to exactly two other segments, except the tips (only one neighbour) and segments at the branching points (connected to three other segments).

To mimic the fluctuating direction of hyphae extension as seen in experiments, we add a small random angle to the orientation of every newly added segments. Since the model does not consider the mechanical properties of the cell wall nor the direction where the hyphae are growing, these random orientations do not affect any processes happening in the cytoplasm. The addition of the new segment happens at the tip or at the branching points once enough macromolecules have been consumed by the polarisome (or TIPOC) (see below).

Connected cell segments provide the space where the processes governing the hyphae growth happen.

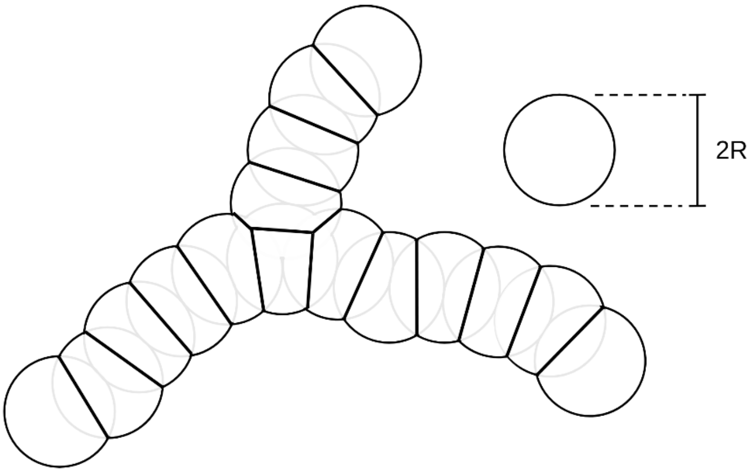

**Figure 1 Geometric representation of the hyphae modeled by a chain of cell segments.** Each cell segment is obtained from non-overlapping parts of intersecting circles with a diameter of 2*R*, the center of which are separated by *R* from the center of the adjacent segments. The image shows 16 segments, one of which is located at a branching point and 3 at the tips.

#### 3.2. Foci

If there are any foci present in a cell segment, the following processes occur:

- **Movement of the foci within the filament.**

The foci can move as a Brownian particle within the cytoplasm (see below), but they can not cross the cell boundary. Foci can, however, transition freely between adjacent cell segments. The cell boundary is modeled by an algorithm that relocates the foci to the inside of the cell if they try to cross the cell wall.

- **Growth**

Our data indicates that foci appear and, at any moment, are present at a variety of sizes. This suggests that there is a certain growth process of foci involved. Here we assume that the foci can grow up to a certain maximum size $r_{max}$:

$$\frac{\mathrm{d}r}{\mathrm{d}t} = k_q[M] \quad \text{if} \quad r < r_{max} \tag{1}$$

Here $r$ is the radius of the foci, $k_q$ is the growth constant that might represent the rate of protein binding to the foci, and $[M]$ is the concentration of macromolecules in the cell segment.

- **Directed movement of foci due to cytoplasm streaming during hyphal growth.**

Each time a new segment is added to the tip of the hyphae, we assume that cytoplasmic material needs to be transported to the newly created space. Thus generated cytoplasmic flow will transport foci in the direction towards the growing tip. The resulting displacement $L$ of a particular foci towards the tip is assumed to be inversely proportional to the distance *d* between that foci and the tip of the hyphae it belongs to (see Figure 2):

$$L = \frac{R}{1 + \frac{d}{E}} \tag{2}$$

Here, $E$ is a length scale that determines how fast the influence of the tip growth decays as we move away from it, and $d$ is the distance from the foci to the tip taking values of $d = \mathbb{N}R$, where $\mathbb{N}$ is a non-negative integer, and $R$ is the cell segment radius. This mechanism ensures that particles closer to the tip move along the growing direction of the hyphae, recapitulating a streaming effect. As shown in Figure 2 and Equation 2, if there are foci located at the tip of the hyphae (at a distance $d_0 = 0$), adding a new cell segment will move that foci at a distance $L_0 = R$ towards the tip, thus relocating it to the newly created space.

This is the main mechanism that models the directed movement of the foci towards the growing ends of the hyphae and ensures that the foci at the very tip will remain at the tip even after a new segment is added. This mechanism was motivated by our experimental observation that some foci are always located at the tip of the hyphae.

- **Creation of new foci after elongation.**

From our experimental measurements, we know that the amount of foci grows linearly with the length of the hyphae with proportionality constant $k_o$ (Figure 4). The constant $k_o$ can be interpreted as the probability of adding new foci to the cytoplasm every time the hyphae are extended by one segment. This way the number of foci is a random variable whose mean value grows linearly with the length of the hyphae with a proportionality factor of $k_o$. New foci are added at random in the middle of one of the first $\mathbb{F}$ segments from the tip.

- **Production of ATP.**

The cell segments that contain foci produce ATP molecules at a rate that is proportional to the total number of foci *P* contained in a given cytoplasmic $i$ segment and their radius $r_k$:

$$\frac{d[ATP]_i}{dt} = \sum_{k=0}^{P} k_{ATP} \cdot r_k^2 \quad (3)$$

Here $k_{ATP}$ is the number of ATP molecules produced per unit of time and per unit of foci radius squared, $r_k$ is the radius of the $k$-th foci, and $P$ is the total number of foci in the $i$−th cell segment. Here we assumed that the ATP production rate increases with the square of the foci radius (foci area).

- **Random movements of foci.**

It has been reported that the bacterial cytoplasm no longer behaves as a simple viscous liquid for particles greater than 30 nm in size, instead, it displays glass-like dynamics and heterogeneity.[6] Additionally, single-particle-tracking experiments have provided evidence for heterogeneously confined diffusion in the cytoplasm of *B. subtilis* bacteria.[7] Thus it is challenging to find a good model for the random motion of large enzymatic complexes as our foci. For the sake of simplicity, we assume that the foci perform Brownian motion with their diffusion constant decreasing with the radius of the foci and viscosity of the solvent according to the Stokes-Einstein relation.[8]

From visual observations of our experimental data from time-lapse fluorescence microscopy, the random movement of the foci seems to be diminished when they are close to the tip as if the tips were limiting the free movement of the foci. Additionally, it has been previously shown that protein diffusion can be hindered on the cell poles in *B. subtilis* [9], while in *E. coli* the protein mobility can be up to 40% slower at the poles compared to the cell center.[9,10] Thus, in the simulation, we assume that the polarisome can interact with the foci by restricting their diffusive step size, and thus slowing down their diffusion. This interaction can be phenomenologically interpreted as an increase in the effective viscosity of the cell segment where the polarisome is located. In simulations, we describe the respective random displacements of the foci according to the overdamped Langevin equation of the Brownian particle leading to displacement increments:

$$\Delta x(\delta t) = \sqrt{2D_{\mathrm{f}}\delta t} \cdot \mathcal{N}(0,1) \tag{4}$$

$$\Delta y(\delta t) = \sqrt{2D_{\mathrm{f}}\delta t} \cdot \mathcal{N}(0,1) \tag{5}$$

$$D_{\mathrm{f}}(r) = \frac{D_0(r)}{(1 + k_{\mathrm{TF}}T)} \tag{6}$$

$$D_0(r) = \frac{k_B\mathcal{T}}{6\pi\eta r} \tag{7}$$

Where $\Delta x$ and $\Delta y$ are the displacement increments in a small time interval $\delta t$ in the $x$ and $y$ directions, respectively, $D_{\mathrm{f}}(r)$ is the effective diffusion coefficient of the foci affected by the polarisome, $D_0$ is free diffusion according to the Stokes-Einstein relation, where $k_B$ is the Boltzmann constant, $\mathcal{T}$ is the temperature, $\eta$ is the effective viscosity of the cytoplasm, $r$ is the foci radius; the presence of the polarisome with mass $T$ in a given cell segment effectively reduces the diffusivity as described by the strength of polarisome-foci interaction $k_{\mathrm{TF}}$, $\mathcal{N}(0, 1)$ is a random number drawn from a normal distribution with mean 0 and standard deviation 1.

Is important to note that for most of the cell segments, the polarisome mass $T$ is zero (see below), and thus the diffusion coefficient of the foci is equal to $D_0$.

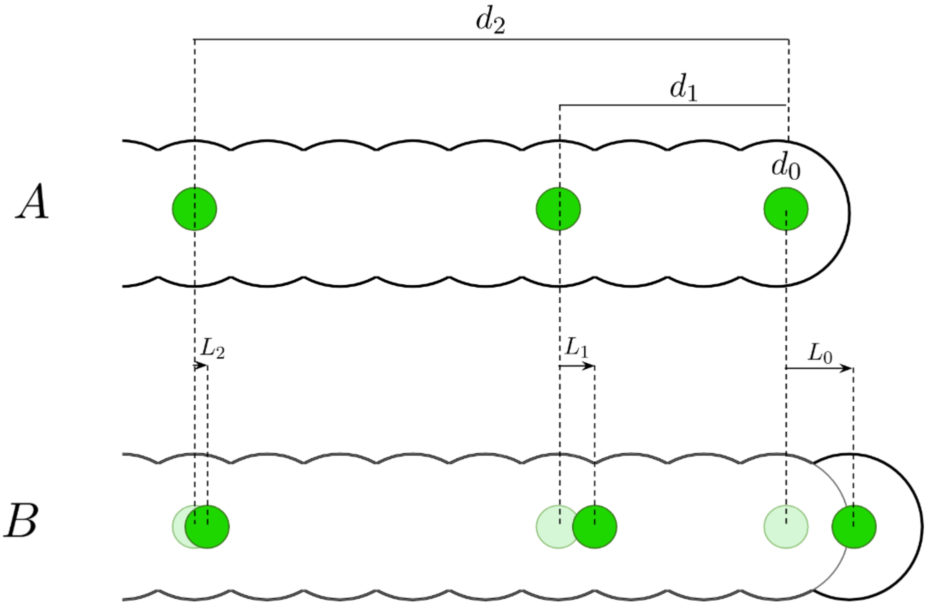

**Figure 2. Directed movement of foci due to cytoplasmic streaming.** A. Before the addition of a new segment, foci are located at a distance $d_0$, $d_1$, and $d_2$ from the tip of the hyphae. B. After the addition of a new segment, the foci are displaced towards the tip of the hyphae by distances $L_0$, $L_1$, and $L_2$, respectively. These displacements are decreasing with the increasing distance from the foci to the tip as described by Eq.(2).

#### 3.3. Polarisome

Polarisome is the ensemble of proteins responsible for adding new cell envelope material to growing regions of the hyphae, building new branches, and elongating the tip of the hyphae.[11] The following rules apply to the polarisome and its associated cell segments:

- **Minimum size required for hyphae elongation and branch initiation.**

Only the cell segments that contain a large enough polarisome with a mass $T > T_{min}$ can elongate or branch. This rule was motivated by the existing models of polarisome splitting in *Streptomyces* spp. that suggest that the polarisome needs to reach a minimum size before it initiates a new branch.[4]

- **Polarisomes consume macromolecules to build cell envelope.**

The polarisome acts as a sink for macromolecules $[M]$ and uses the total amount of “consumed” macromolecules to accumulate new cell envelope material $C$ on the cytoplasm segment where it is located (see below). Once enough of the cell envelope material has been accumulated and a certain threshold $C_{max}$ is reached, a new segment is added to the hyphae. The hyphae are extended by one segment if the polarisome is at the tip, or a new branch is created if the polarisome is located at a branching point. The rate at which a cell segment builds new cell envelope material is the same as the rate at which the segment’s polarisome consumes macromolecules. These rates depend on the size of the polarisome and the concentration of macromolecules:

$$\frac{\mathrm{d}C}{\mathrm{d}t} = k_u T[M] \tag{8}$$

where $C$ is the amount of cell envelope material, $[M]$ is the amount of macromolecules, $T$ is the mass of the polarisome and $k_u$ is the macromolecule consumption rate constant associated with the polarisome.

- **Polarisome growth.**

Previous models of polarisome splitting have included the binding rate of proteins to the polarisome as a parameter that describes the growth of the polarisome.[12] Here we assume that the polarisome grows at a rate that is proportional to the amount of freely diffusing macromolecules available in the cell segment where the polarisome is located. This way, the polarisome will grow faster in regions where there are more macromolecules available and the probability of new proteins binding to the polarisome is higher:

$$\frac{\mathrm{d}T}{\mathrm{d}t} = k_T[M] \qquad if \quad T < T_{max} \qquad (9)$$

where $k_T$ is the polarisome growth rate constant that represents the binding rate of proteins to the polarisome.

- **Splitting of polarisome.**

Once the polarisome has reached its maximum allowed mass $T_{max}$, it will asymmetrically split to create new branches, leaving one smaller polarisome particle behind at the branching point and moving the other larger part of the polarisome to the growing tip of the hyphae. The proportion $P_T$ of polarisome that is left behind at the branching point is a random variable sampled from a uniform distribution. This mechanism was motivated by previous models of polarisome splitting.[12]

### 4. Cellular concentrations modeled by continuous fields

#### 4.1. ATP

The model describes ATP production, consumption, and diffusion within the hyphae. The following processes occur:

- **Production of ATP.**

ATP molecules are locally produced by the cytoplasmic segments that contain foci. The rate of ATP production is proportional to the total number of LpdA-containing foci *P* in the respective cytoplasm segment, see Eq.3.

- **Consumption.**

It is known that synthesis of proteins and lipids is the cellular processes that consume the highest proportion of the cellular energy budget[5], and protein synthesis and cell envelope synthesis are carried out near the tips of the hyphae.[3,4,13] Therefore, we assume that the consumption rate of ATP is higher in the segments that are closer to the tips. Even though we expect the ATP consumption rate to be lower in the segments that are further away from the tips, it is never zero in any segment due to the housekeeping processes. The ATP consumption by these two processes (cell envelope synthesis and housekeeping) is modeled by the following equation:

$$\frac{\mathrm{d}[ATP]}{\mathrm{d}t} = -\left(\frac{k_c}{1 + d/E_a} + k_h\right)[ATP] \quad (10)$$

Where $E_a$ is the length scale describing the decay of tip influence (analogous to $E$ in equation 2), $k_c$ is the ATP consumption rate constant associated with the synthesis of macromolecules, $[ATP]$ is concentration of ATP in the cytoplasmic segment, $d$ is the distance of the cell segment from the tip of the hyphae, and $k_h$ is the ATP consumption rate constant associated with housekeeping processes.

- **Diffusion.**

ATP diffuses between adjacent cell segments following the Fick's law of diffusion. We model this process at the level of segment discretization. For a typical cell segment $i$ that has two other segments connected to it ($i$ − 1 and $i$ + 1), the diffusion of ATP is described by the following equation:

$$\Delta[ATP]_i = \frac{D_A \Delta t}{\Delta x^2}\left([ATP]_{i-1} - 2[ATP]_i + [ATP]_{i+1}\right) \quad (11)$$

Where $\Delta[ATP]_i$ is the change of ATP content in the cell segment $i$ in a time step $\Delta t$, $D_A$ is the diffusion coefficient of ATP molecule, $\Delta x = R$ is the distance between the centers of the adjacent cell segments.

For a cell segment $i$ that has only one other segment $i$ + 1 connected to it (i.e. at the tip), the diffusion of ATP is described by the following equation:

$$\Delta[ATP]_i = \frac{D_A \Delta t}{\Delta x^2}\left([ATP]_{i+1} - [ATP]_i\right) \quad (12)$$

For a cell segment $i$ that is located at a branching point and has three other segments connected to it ($j$, $k$, and $l$), the diffusion of ATP is described by the following equation:

$$\Delta[ATP]_i = \frac{D_A \Delta t}{\Delta x^2}\left([ATP]_j + [ATP]_k + [ATP]_l - 3[ATP]_i\right) \quad (13)$$

#### 4.2. Macromolecules

Macromolecules in the model is a very broad term that represents all the building blocks for cellular structures that are part of the cell envelope and cytoplasm.

Macromolecules can be produced, consumed, and diffused according to the following rules:

- **Production.**

Macromolecules are synthesized using ATP. They are produced at a higher rate near the tips of the hyphae, where RNA translation and protein synthesis are more active.[3] The production rate of macromolecules is therefore described by the following equation:

$$k_a \frac{\mathrm{d}[M]}{\mathrm{d}t} = \frac{k_c[ATP]}{1 + d/E_a} \qquad (14)$$

where $k_a$ is the number of ATP molecules that are required to produce one macromolecule, and $k_c$ is the ATP consumption rate constant associated with the synthesis of macromolecules (same as in Eq. 10).

- **Consumption.**

All cell segments are thought to represent sinks for macromolecules, where they are either consumed by the polarisome or degraded in the natural turnover of the cell.

The consumption rate is proportional to the size of the polarisome and the concentration of macromolecules:

$$-\frac{\mathrm{d}[M]}{\mathrm{d}t} = (k_u T + k_b)[M] \qquad (15)$$

Here, $k_u$ is the macromolecules consumption rate constant associated with the polarisome (same as in Eq. 8), and $k_b$ is the basal macromolecules consumption rate constant that is present in all cell segments and represents macromolecules turnover.

- **Diffusion.**

Like ATP, macromolecules diffuse between cell segments following the Fick's law of diffusion (equations 11, 12, and 13), where we need to substitute $[M]$ instead of $[ATP]$, and the respective diffusion constant of macromolecules $D_M$.

- **New cytosolic segment addition.**

As soon as in the segment where polarisome is located (tip or a branching point) the amount of consumed macromolecules crosses a threshold $C > C_{max}$, a new empty cell segment is added at the tip (branching) point.

#### 5. Parameters

The model parameters can be grouped into several categories based on their role in the simulations. Here we provide a detailed overview of these parameters and their justification. The summary of all model parameters, their descriptions, and values can be found in Table 1.

##### 5.1. Spatial scales

- **Radius of a cell segment** ($R$ = 0.5 µm): Based on previous measurements of *Streptomyces* hyphae and our experimental observations.[14]
- **Distance between segment centers** ($\Delta x$ = 0.5 µm): Chosen to match the cytoplasm radius for optimal spatial discretization.
- **Tip influence decay length for streaming of foci** ($E$ = 1 µm): Fit parameter adjusted to match the resulting uniform foci distribution along the hyphae.
- **Tip influence decay length for ATP consumption** ($E_a$ = 4.5 µm): fitted parameter adjusted to align with ATP consumption rates observed in *E. coli* during the exponential growth and stationary phases.[15,16] In *E. coli*, these rates are $6.4\times10^6$ ATP/s during exponential growth and $0.4\times10^6$ ATP/s during the stationary phase.[15] For *Streptomyces* hyphae, we estimate ATP consumption at the tip to be similar to *E. coli* in exponential growth, while segments farther from the tip are expected to have rates comparable to the stationary phase.

##### 5.2. Diffusion constants

- **ATP diffusion coefficient** ($D_A$ = 100 µm$^2$/s): Due to the glass-like properties of the cytoplasm[6], the perceived cytoplasmic viscosity has been shown to be dependent on the size of the molecule that is diffusing. For small molecules the cytoplasm viscosity has values between 0.006 Pa·s and 0.001 Pa·s; which is not far from the viscosity of water. Particularly, for the BCECF molecule (~800 Da), the cytoplasmic diffusion coefficient is approximately four times slower than in water ($D_{cyto}/D_{water} \sim 0.25$).[17,18] Here we will assume that ATP (~ 500 Da) diffusion coefficient in the cytoplasm is reduced by a factor of ~ 4 compared to the diffusion coefficient of ATP in water. The diffusion coefficient in water has previously been reported to be 387 µm$^2$/s.[19]

This means that we can estimate the cytoplasmic diffusion coefficient of ATP to be approximately 100 μm²/s.

- **Macromolecule diffusion coefficient** ($D_M$ = 1.56 μm²/s): The cytoplasm contains a diverse mixture of macromolecules with varying sizes, which makes it difficult to get a single diffusion coefficient for all these molecules, however, we can use an approximated value based on the typical size of the particles that are contained in the cytoplasm. Some studies have estimated that the cytoplasm of *E. coli* contains approximately $3\times10^6$ globular particles (macromolecules), each with an average radius of 2.5 nm.[20] If we use the mean cytoplasmic viscosity of *B. subtilis* of 0.0549 Pa·s [9] and the Stokes-Einstein relation for spherical particles of radius 2.5 nm, we can estimate the diffusion coefficient of these particles to be 1.56 μm²/*s*.

#### 5.3. Foci parameters

- **Foci radius** ($r$ = 6.6 nm to 150 nm): From our experimental measurements, the apparent radius of the foci is $r_a \approx 0.25$ μm which is the same order as the Airy disk of a point-like light source, indicating that the aggregates are rather small.

  To simulate the Brownian motion of the foci according to the Stokes-Einstein relation and their growth, we need to assign a certain initial radius $r > 0$ to them. Even though the structure of the PDH-ODH complex containing LpdA is not fully known, the minimal hybrid PDH-ODH complex is estimated to be about 1 MDa.[1] Assuming that the PDH-ODH complex forms a sphere with a partial specific volume of 0.73 cm³/g, we can estimate its minimum radius with the following formula:

$$R_{min} = 0.66 \cdot m^{1/3} \tag{16}$$

  where $R_{min}$ is the radius of the sphere in nanometers and *m* is the mass of the complex in Dalton.[21,22] This gives us an estimate of the minimum radius of the PDH-ODH complex to be 6.6 nm. Since enzymatic megacomplexes are usually 30-300 nm in diameter[6], we can assume that the radius of the foci to be bounded between 6.6 nm and 150 nm. In the simulations, we have chosen 6.6 to be the radius of the nucleation sites where new foci are created and 150 nm to be the maximum radius $r_{max}$ that the foci can reach.

- **Foci growth constant** ($k_q$ = 2×10$^{-11}$ µm/(s·molecule)): This is a fit parameter adjusted so that the foci can reach its maximum size in a period in the order of minutes.
- **Region for addition of new foci:** $d_i = \mathbb{F}R$, where $\mathbb{F}$ is an integer and $\mathbb{F} \sim U(0, 7)$ Here, $d_i$ is the distance from the tip of the hyphae where new foci are added, and $R$ is the radius of the cell segment, $\mathbb{F}$ is a random integer sampled from a uniform distribution between 0 and 7. This parameter is based on experimental observations and literature data that suggest that DNA transcription is spatially constrained at around 2 µm behind the extending tip.[3] If we assume that protein synthesis happens near the transcription site, we can infer that new foci are being added around this region.
- **Probability of adding foci after tip extension** ($k_o$ = 0.34): Fit parameter adjusted to match the experimentally observed amount of foci per unit of length of hyphae (Figure 4).
- **Interaction strength between foci and polarisome** ($k_{FT}$ = 20 fg$^{-1}$): Fit parameter adjusted to mimic foci confinement near the tip due to molecular crowding and polarisome-foci interaction.
- **Cytoplasm effective viscosity** ($\eta$ = 6 Pa·s): The perceived intracellular viscosity has been shown to be dependent on the size of the molecule that is diffusing, with larger molecules perceiving the cytoplasm not as a simple viscose liquid, but as a glass-like medium.[6]
  Some authors have reported the mean viscosity of the *B. subtilis* bacterial cytoplasm to be approximately 0.0549 Pa·s (about 50 times the viscosity of water).[9] Given the large size of the foci, we estimate for LpdA-foci an effective viscosity to be higher than the average viscosity for *B. subtilis*. In our simulations, we selected a value of 6 Pa·s as a first approximation for the effective viscosity of LpdA-containing foci diffusing in the cytoplasm.
- **Temperature of the cytoplasm** ($\mathcal{T}$ = 293 K = 19.85 °C)

#### 5.4. Polarisome parameters

- **Mass threshold for elongation/branching** ($T_{min}$ = 0.5 fg): According to previous models of tip splitting, there is a minimum size that the polarisome must reach before it extends the hyphae or create a new branch.[12] We propose this value as a first approximation.

- **Mass threshold for splitting** ($T_{max}$ = 1 fg): Similar to $T_{min}$, there is a certain size that the polarisome must reach before it splits.[12] We propose this value as a first approximation.
- **Polarisome growth constant** ($k_T = 4\times10^{-12}$ fg/(s·molecule)): Growth rate of the polarisome. This is a fit parameter adjusted to achieve observed tip extension rate and branching frequency. If this value is too high, the polarisome will grow too fast and will reach the splitting size too quickly, producing new branches with a high frequency. If the value is too low, the polarisome will split infrequently, producing fewer branching points.
- **Polarisome splitting ratio** ($P_T \in \mathbb{R}$ and $P_T \sim$ U(0, 0.5)): Previous models suggest that the polariosme splits asymmetrically.[12]

#### 5.5. Metabolic rates

- **ATP production rate** ($k_{ATP} = 1.45\times10^8$ molecules/(s·µm$^2$)): This is a fit parameter that was adjusted to match the ATP consumption rates and cellular ATP concentrations reported in the literature.[15,16,23–25]
- **ATP consumption rate for macromolecule synthesis** ($k_c$ = 1.94 s$^{-1}$): Estimate based on literature data on cellular ATP consumption rates and ATP concentration.[15,16,23–25]
- **ATP consumption rate for housekeeping processes** ($k_h$ = 0.9 s$^{-1}$): Estimate based on literature data on cellular ATP consumption rates and ATP concentration.[15,16,23–25]
- **Macromolecule consumption rate for cell envelope synthesis** ($k_u = 3\times10^{-4}$ s$^{-1}$ fg$^{-1}$): Fit parameter adjusted to match observed tip extension rates previously reported in the literature[26] and keep amount of macromolecules in the cell segment close to $3\times10^6$.[20]
- **Macromolecule decay rate** ($k_b = 9\times10^{-5}$ s$^{-1}$): Fit parameter adjusted to keep the amount of macromolecules in the cell segment close to $3\times10^6$.[20] If this parameter is too low, the macromolecules will accumulate in the cell segment.

#### 5.6. Conversion factors

- **ATP per macromolecule** ($k_a$ = 1×10$^3$ ATP molecules/macromolecule): Fit parameter adjusted to match expected hyphae growth rate and also based on estimates on the average energy cost of protein synthesis.
- **Macromolecules required for creating a new cell segment** ($C_{max}$ = 10×10$^6$ macromolecules/segment): Fit parameter adjusted to match observed tip extension rates.

Future work should focus on experimental validation of fit parameters and refinement of literature-derived values for the specific context of *Streptomyces* spp. growth.

*Table 1 Summary of all model parameters, their descriptions, and values. Parameters are grouped by their functional categories.*

| Parameter | Description | Value |
|---|---|---|
| | **Spatial Scales** | |
| $R$ | Radius of cell segment | 0.5 µm |
| $\Delta x$ | Distance between segment centers | 0.5 µm |
| $E$ | Tip influence decay length for streaming of foci | 1 µm |
| $E_a$ | Tip influence decay length for ATP consumption | 4.5 µm |
| | **Diffusion Constants** | |
| $D_A$ | ATP diffusion coefficient | 100 µm$^2$/s |
| $D_M$ | Macromolecule diffusion coefficient | 1.56 µm$^2$/s |
| | **Foci Parameters** | |
| $r$ | Foci radius range | 6.6–200 nm |
| $k_q$ | Foci growth constant | 2×10$^{-11}$ µm/(s·molecule) |
| $k_o$ | Probability of adding foci after tip extension | 0.34 |
| $k_{FT}$ | Interaction strength between foci and polarisome | 20 fg$^{-1}$ |
| $\eta$ | Cytoplasm viscosity for foci | 6 Pa·s |

*Continued on next page*

| | | |
|---|---|---|
| $\mathcal{T}$ | Temperature of cytoplasm | 293 K |
| | **Polarisome Parameters** | |
| $T_{min}$ | Mass threshold for elongation/branching | 0.5 fg |
| $T_{max}$ | Mass threshold for splitting | 1 fg |
| $k_T$ | Polarisome Growth constant | $4\times10^{-12}$ fg/(s·molecule) |
| $P_T$ | Polarisome splitting ratio | U(0, 0.5) |
| | **Metabolic Rates** | |
| $k_{ATP}$ | ATP production rate | $1.45\times10^{8}$ molecules/(s·$\mu m^2$) |
| $k_C$ | ATP consumption rate for macromolecule synthesis | 1.94 $s^{-1}$ |
| $k_h$ | ATP consumption rate for housekeeping processes | 0.9 $s^{-1}$ |
| $k_u$ | Macromolecule consumption rate for cell envelope synthesis | $3\times10^{-4}$ $s^{-1}\cdot fg^{-1}$ |
| $k_b$ | Macromolecule decay rate | $9\times10^{-5}$ $s^{-1}$ |
| | **Conversion Factors** | |
| $k_a$ | ATP per macromolecule | $1\times10^{3}$ ATP/macromolecule |
| $C_{max}$ | Macromolecules required for new cell segment | $10\times10^{6}$ macromolecules/segment |

### 6. Results

We simulated 20 realizations of the model with the duration corresponding to 8 hours of growth and registered the evolution of the total length of the hyphae in 5-second intervals. These simulation results were compared to the experiments on growing *Streptomyces coelicolor* (Experiment 1) and *Streptomyces venezuelae* (Experiment 2) where we measured the total length of the hyphae and the number of foci at intervals of 10 minutes during 10 hours in two replicate experiments.

#### 6.1. Growth rates

The growth curves from the experiments and simulations are shown in Figure 3.

The simulated growth rate falls between the two experimental measurements, indicating good agreement between the model and experimental observations. The similar slopes between experimental and simulated data suggest that the model accurately captures the exponential growth behavior of *Streptomyces* hyphae. We should note that there is a phase of slow growth (~ 5 h) due to the germination of spores in the experiments that are not modeled in the computational model. This discrepancy is reflected in the difference between the length of the hyphae in simulations compared to the length of the hyphae in the experiments.

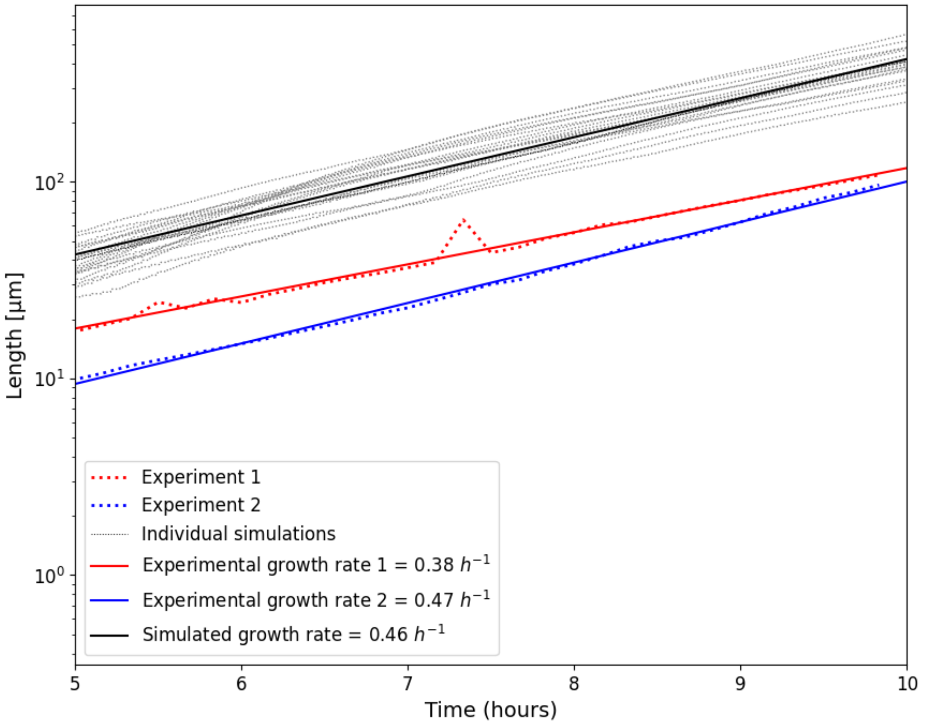

**Figure 3. Comparison of experimental and simulated growth rates.** The graph shows both experimental and simulated growth data for *Streptomyces* hyphal growth. Two experimental datasets (red and blue dotted lines) are shown with their corresponding linear regressions, yielding growth rates of 0.38 $h^{-1}$ and 0.47 $h^{-1}$ respectively. The datasets from simulations (gray lines) are shown with their average growth rate represented by a black line showing a growth rate of 0.46 $h^{-1}$.

#### 6.2. Foci count per unit of length

We also measured the amount of foci per unit of length of the hyphae in the experiments to tune the value of $k_o$ in the model. In Figure 4, we show the total count of foci from the two replicates of *Streptomyces* spp. growth experiments and also the total count of foci per unit of length from 20 simulation runs. This close agreement between experimental and simulated foci densities validates our interpretation of the parameter $k_o$ as the probability of foci addition after tip extension (see 5.3.).

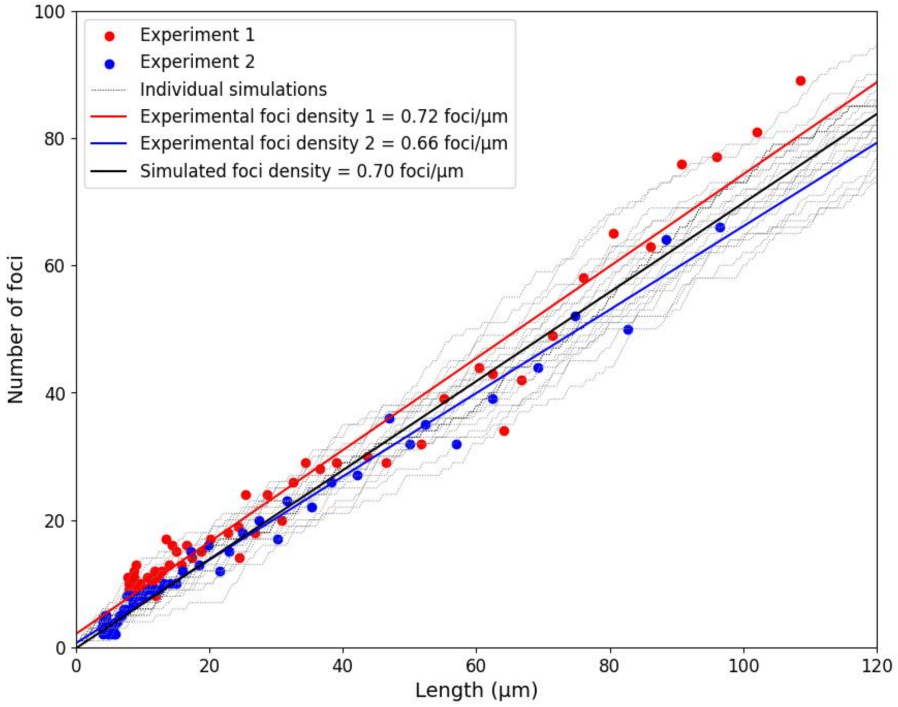

**Figure 4. Comparison of experimental and simulated foci counts.** The graph shows both experimental data and simulation results for the number of foci as a function of hyphal length. Experimental data points are shown from two replicates (Experiment 1 (*S. coelicolor*) in red and Experiment 2 (*S. venezuelae*) in blue), with their corresponding linear regressions yielding foci densities of 0.72 and 0.66 foci per micrometer, respectively. The simulation results from 20 individual runs (shown as thin gray lines) demonstrate the stochasticity of foci formation, with their average foci density represented by a black line (0.70 foci per micrometer).

#### 6.3. Spatial heterogeneity in metabolic activity

Our simulations reveal distinct spatial heterogeneity in metabolic activity throughout *Streptomyces* hyphae (Figure 5). This heterogeneity likely arises from stochastic processes such as foci addition and movement, polarisome splitting, and elevated ATP consumption near hyphal tips compared to other regions (Figure 5, right panel).

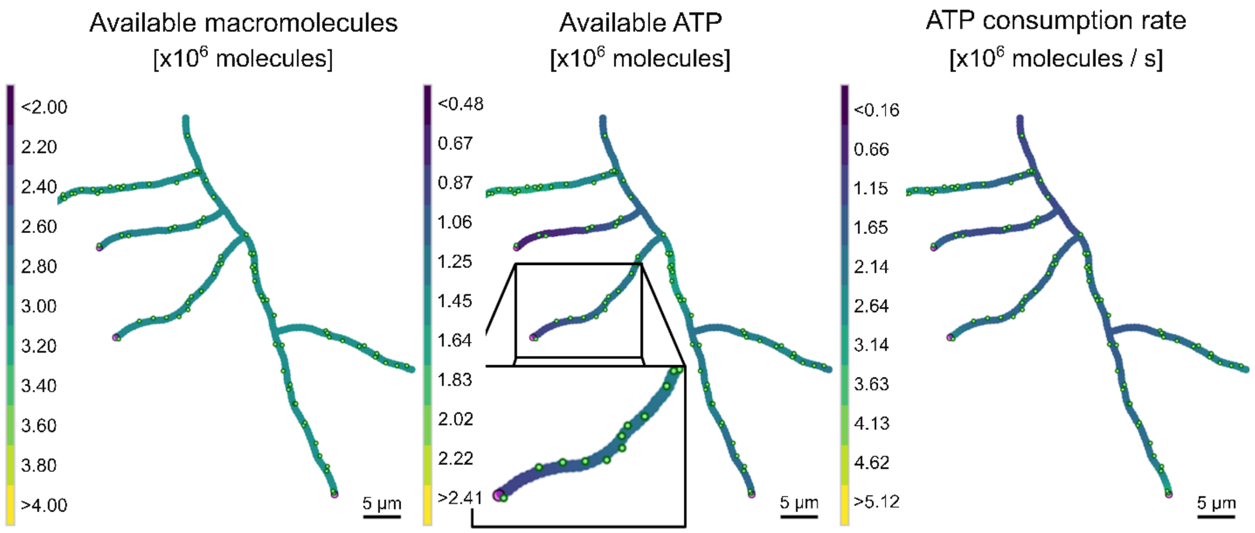

**Figure 5. Macromolecules concentration, ATP concentration, and consumption rate.** The concentration of freely diffusing macromolecules is shown on the left panel. The panel at the center shows the count of ATP molecules contained in each individual cell segment. The color scale represents the concentration of macromolecules and ATP in $10^6$ molecules per cytoplasm segment. The right panel shows the rate at which ATP is consumed in each cytoplasm segment. The color scale represents the consumption rate in $10^6$ molecules per second per cytoplasm segment. The LpdA-containing foci are represented by the green dots, and the polarisomes are represented by the purple dots at the tips of the hyphae.

Figure 5 shows typical model outputs under normal conditions, replicating the usual growth pattern of *Streptomyces* species. It compares the concentrations of freely diffusing macromolecules and ATP molecules as well as the ATP consumption rate, highlighting their dynamic relationship throughout the hyphae.

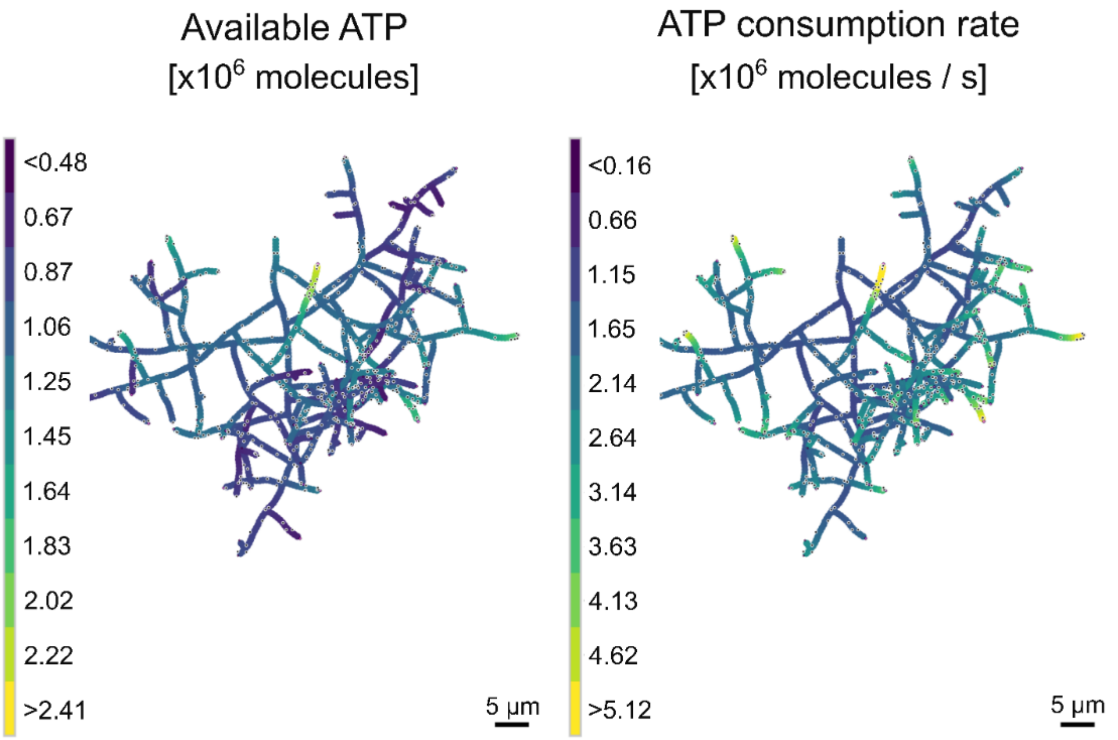

**Figure 6. Hyperbranching simulation.** The image illustrates the increased branching frequency when the model is simulated with a lower value for polarisome splitting size.

#### 6.4. Simulating hyperbranching

By adjusting parameter settings, the model can also simulate phenotypic variations like hyperbranching (Figure 6). This simulation of the hyperbranching phenotype, characterized by increased branching frequency, can be induced by decreasing the maximal size at which polarisomes start to split. This causes more frequent splitting events, each one generating a new branch.

### 7. Numerical implementation

The numerical implementation of simulations is facilitated by noticing scale separation of the two different time scales. The overall structure follows a hierarchical organization where faster processes are nested within slower ones to ensure numerical stability and computational efficiency.

The simulation employs two time-scales to handle processes that occur at different rates:

- **Fast processes** (Numerical time step $\Delta t$ = 0.825 ms):
  Chemical reactions and diffusion are simulated at this time scale. This includes simulating the production, consumption, and diffusion of ATP and macromolecules within each cytoplasm segment. The reaction-diffusion time step $\Delta t$ is chosen to satisfy the Courant-Friedrichs-Lewy (CFL) condition for numerical stability[27]:

$$\Delta t \leq \frac{(\Delta x)^2}{2D_A w} \qquad (16)$$

  where $w$ is the number of dimensions in the system, and $D_A$ is the highest diffusion coefficient in the system, which is the diffusion coefficient of ATP. For computational efficiency, one aims to choose the largest possible value of $\Delta t$ that satisfies the CFL condition, which is $\Delta t = (\Delta x)^2/(2D_A w)$.
  The filament without branches could be considered a one-dimensional system, and thus, one could choose $w$ = 1. However, due to the branching points, a value of $w$ = 1 makes the simulation unstable as soon as a new branch is created. A value of $w$ = 2 would be adequate for a simulation in which the substances diffuse in a 2D plane. Even though we could have chosen $w$ = 2 to ensure numerical stability, the intermediate value of $w$ = 1.5 was optimal to ensure both numerical stability and computational efficiency.

- **Intermediate processes** (numerical time step $\delta t$ = 5 s):
  This time step is used to iterate the Brownian motion of foci, and the growth of polarisome. Additionally, at this time scale, the simulation checks for conditions that trigger events such as the addition of new cell segments, creation of new branches, and addition of new foci.

This implementation with two time-scales ensures that each process is simulated at an appropriate temporal resolution while maintaining numerical stability and computational efficiency. The hierarchical organization allows for accurate representation of both fast molecular processes and slower morphological changes in the growing *Streptomyces* colony.

### 8. Conclusion

We developed an agent-based computational model to explore the possible roles of LpdA-containing foci in the growth and branching behavior of *Streptomyces* hyphae. The model incorporates key cellular components, including cell segments, polarisomes, LpdA-containing foci, ATP, and macromolecules.

Our model shows that *Streptomyces* hyphae likely exhibit spatial heterogeneity of metabolic activity along the filament. This phenomenon is attributed to the high consumption rates of ATP and macromolecules in the regions near the tips and the local production of ATP by the foci.

The model also demonstrates its versatility by simulating different growth patterns, including normal growth and hyperbranching phenotypes. By adjusting parameters such as the polarisome splitting size, we can induce increased branching frequency, providing insights into how modifications to the model parameters can lead to alterations to the morphology and cytoplasm concentrations in the hyphae.

Future work should focus on refining the model by incorporating realistic parameters for diffusivities and reaction rates. Importantly, it would be also possible to incorporate (to be studied) the biophysical mechanism of foci formation.
